# Changes in gene expression during female reproductive development in a colour polymorphic insect

**DOI:** 10.1101/714048

**Authors:** B. Willink, M. C. Duryea, C. Wheat, E. I. Svensson

## Abstract

Pleiotropy (multiple phenotypic effects of single genes) and epistasis (gene interaction) have key roles in the development of complex phenotypes, especially in polymorphic taxa. The development of discrete and heritable phenotypic polymorphisms often emerges from major-effect genes that interact with other loci and have pleiotropic effects on multiple traits. We quantified gene expression changes during ontogenetic colour development in a polymorphic insect (damselfly: *Ischnura elegans*), with three heritable female morphs, one being a male mimic. This female colour polymorphism is maintained by male mating harassment and sexual conflict. Using transcriptome sequencing and *de novo* assembly, we demonstrate that all three morphs downregulate gene expression during early colour development. The morphs become increasingly differentiated during sexual maturation and when developing adult colouration. These different ontogenetic trajectories arise because the male-mimic shows accelerated (heterochronic) development, compared to the other female morphs. Many loci with regulatory functions in reproductive development are differentially regulated in the male-mimic, including upstream and downstream regulators of ecdysone signalling and transcription factors potentially influencing sexual differentiation. Our results suggest that long-term sexual conflict does not only maintain this polymorphism, but has also modulated the evolution of gene expression profiles during colour development of these sympatric female morphs.

## Introduction

The role of epistasis (gene interaction) in shaping phenotypic variation and covariation has been discussed since the early days of the Modern Synthesis (Fisher 1930; Wright 1930; 1931), and still remains a contentious subject (Hill et al. 2008; Shao et al. 2008; Huang et al. 2012; Hemani et al. 2014; Mackay 2014; Wood et al. 2014). Epistasis occurs when the phenotypic effect of a gene or genetic variant depends on the genotype of other loci. Major-effect loci affect multiple traits by operating on top of gene regulatory networks (Tyler et al. 2009). Allelic variants in major-effect loci can therefore cause extensive phenotypic differentiation by interacting epistatically with downstream genes in modular regulatory networks (Monteiro and Podlaha 2009).

Colour-polymorphic taxa provide interesting model systems for investigating both the scope and mechanisms of such pleiotropy of master-regulatory loci via gene interactions. Heritable colour morphs typically differ in multiple physiological, developmental and life-history traits (McKinnon and Pierotti 2010), while having a relatively simple genetic basis of one or a few loci with restricted recombination in between them (Joron et al. 2006; Lamichhaney et al. 2016; Lindtke et al. 2017; Andrade et al. 2019). The genetic basis and molecular mechanisms behind the phenotypic differences of such heritable colour morphs can therefore reveal direct or indirect interactions between colour-morph loci and other loci influencing suites of phenotypic traits. While such widespread pleiotropic effects of colour-morph loci are increasingly being recognized (Gangoso et al. 2016; Dijkstra et al. 2017; Toomey et al. 2018; Woronik et al. 2019), we know little about the gene-regulatory effects of colour-morph loci, during the critical developmental periods when extensive phenotypic differences between morphs typically emerge.

Phenotypic evolution in animals has many times resulted from regulatory changes in the timing, location and volume of interactions among gene products (Gerhart and Kirschner 2007). Many evolutionary novelties have arisen through a shift in the relative timing or rate of gene expression (e.g. Gomez et al. 2008; Albertson et al. 2010; Gunter et al. 2014), rather than due to the disruption of previous interactions or through the formation of entirely novel networks. This phenomenon, known as heterochrony was historically considered to be a source of novel phenotypic variation between independently evolving lineages (Gould 1977). However, variation in the timing of developmental events can also be important at the intraspecific level (Linksvayer and Wade 2005). Heterochrony can produce large phenotypic change with only small genetic change (West-Eberhard 2003). Heterochrony may therefore contribute to pervasive phenotypic differences between intraspecific colour morphs, if genetic variation in the morph-determining locus controls the timing or relative order of developmental events within morphs.

A small but increasing body of evidence from several colour polymorphic taxa suggests that genetic control over regulatory networks can produce substantial intraspecific phenotypic variation (Kunte et al. 2014; Timmermans et al. 2014; Yassin et al. 2016; Takahashi et al. 2018). Such gene regulatory networks may modify both the identity of interacting genes and the relative timing of conserved interactions (e.g. Pham et al. 2017). Developmental characterisation of gene expression profiles of colour polymorphic taxa is a first step to assess how variation in regulatory targets and in the timing of regulatory events cause extensive phenotypic differences between morphs. By investigating direct or indirect interactions between colour-morph loci and other loci during development, we can increase our understanding of how epistasis and pleiotropy shape different morphs within species.

Discrete sympatric morphs can emerge as an evolutionary outcome of sexual conflict over mating (Gavrilets and Waxman 2002). For instance, in several insect taxa, female-limited phenotypic polymorphisms have evolved, and their maintenance has been linked to sexual conflict over mating (Svensson et al. 2005; 2009; Reinhardt et al. 2007; Karlsson et al. 2013; Iversen et al. 2019). More recently, such female-limited polymorphisms have also been described in a few vertebrate species, and appear to be promoted by male mating harassment and sexual conflict (Lee et al. 2019; Moon and Kamath 2019). Here, we investigated how changes in gene expression during adult development (i.e. after the last moult) differ among heritable female morphs in the trimorphic common bluetail damselfly (*Ischnura elegans*). Several damselfly species in this genus, including *I. elegans*, express such female-limited colour polymorphisms that are maintained by frequency-dependent sexual conflict, and in which common morphs suffer from reduced fitness due to excessive male mating harassment (Svensson et al. 2005; Gosden and Svensson 2009; Takahashi et al. 2010; Le Rouzic et al. 2015, Gering 2017).

Two of the female morphs in *I. elegans* differ phenotypically from males in their sexually mature colouration, while the third morph is considered to be a male-mimic, with a colour pattern strikingly similar to that of males (Svensson et al. 2009). Male-mimicking females also differ from the other two morphs in their ontogenetic trajectories of colour change and in several other phenotypic traits, including fecundity (Svensson and Abbott 2005; Willink et al. 2019), mating rates (Gosden and Svensson 2009), parasite resistance and tolerance (Willink and Svensson 2017), cold tolerance (Lancaster et al. 2017) and adult development times (Svensson et al. 2020). Some of these phenotypic differences between the morphs are likely to be adaptive and a result of correlational selection (Svensson 2017). Proximately, morph differences may arise because the colour locus (or a set of tightly linked loci) directly or indirectly regulates other loci affecting co-selected traits. Such regulatory interactions can be investigated by comparing gene expression profiles during ontogenetic development of the female morphs. Quantitative gene expression studies of polymorphic taxa are therefore useful study systems for biologists who are interested in the evolution of gene regulatory networks, and how they are influenced by sexual conflict and sexually antagonistic selection (Hill et al. 2019).

We compared gene expression profiles during ontogeny and colour development of the three female morphs in *I. elegans*. The transcriptional differences that we describe and quantify in this study reveal pleiotropic effects of the colour locus and illustrate the evolutionary outcome of the long-term sexual conflict and the resulting frequency-dependent selection that has shaped this female polymorphism. Specifically, we compare gene expression profiles of the male-mimicking female morph and the two female morphs that are distinctly different from males in their sexually mature colour patterns (Fig. 1a). We address several specific questions about the developmental regulation of gene expression during colour development and how it differs between female morphs. First, at which point(s) in colour development do female morphs diverge in their gene expression profiles? Second, which physiological processes characterise colour development across the female morphs of *I. elegans?* Third, which regulatory genes could cause these differences between the morphs? Finally, are differences in fecundity between morphs due to heterochrony of sexual maturation with respect to colour development?

**Figure 1.**
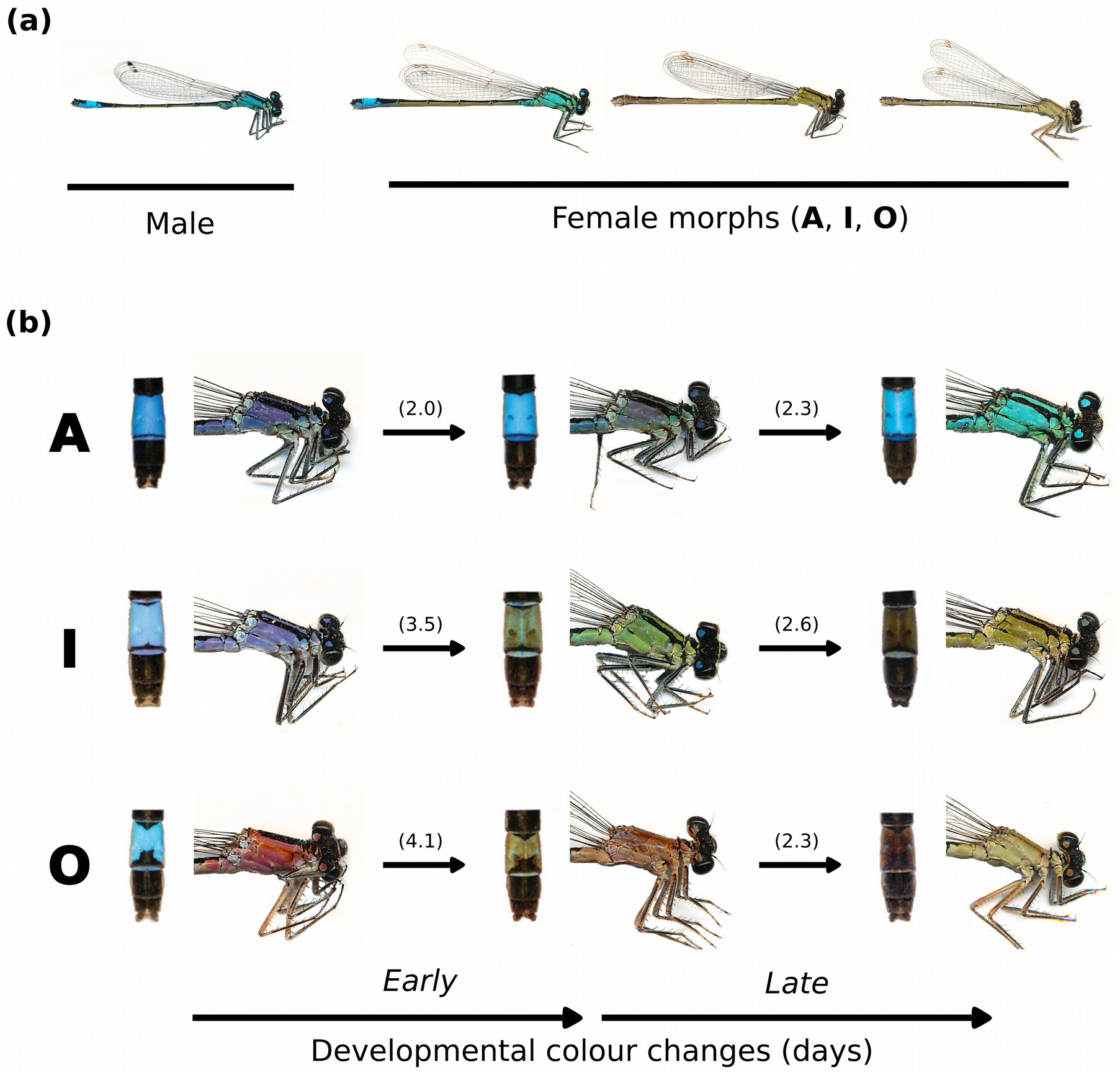
Heritable colour polymorphism and ontogenetic colour changes in females of the common bluetail damselfly (*Ischnura elegans*). **(a)** Male and the three female-limited colour morphs of *I. elegans* in their final, sexually mature developmental phase. The colour pattern of sexually mature A-females is remarkably similar to that of males, and hence they are considered to be male mimics. **(b)** Ontogenetic colour changes in the distal abdomen segments (left) and the thorax (right) of females of the three colour morphs in *I. elegans*. The columns correspond to females in the *immature*, *transitional* and *mature* developmental colour phases. The rows represent colour changes within each heritable morph. The numbers in parenthesis give the mean number of days for each colour transition, when females were kept in large outdoor mesocosm enclosures from their immature phase until sexual maturity (see Fig. S1).

Apart from gaining a better understanding of the molecular mechanisms underlying the female colour polymorphism in *I. elegans* and other polymorphic damselflies, the answers to the questions above should also be of general interest for the genomic consequences and evolutionary outcomes of various forms of sexual conflict (Rowe et al. 2019). With a few recent notable exceptions (Barson et al. 2015; Cheng and Kirkpatrick 2016; Dutoit et al. 2018), our current empirical knowledge about the genomic signature and consequences of sexual conflict is extremely limited (Rowe et al. 2019). The results in the present study suggest that not only can sexual conflict maintain sexually selected polymorphisms through balancing selection, but such conflict can also shape patterns of pleiotropy, epistasis, gene expression profiles and gene regulatory networks associated with such polymorphisms (cf. Hill et al. 2019).

## Methods

### Study system

Females of the common bluetail damselfly (*Ischnura elegans*) occur in three heritable colour morphs whereas males are monomorphic (Fig. 1a). Female morphs differ in both their sexually mature (final) colour patterns as well as in the qualitative colour changes that are expressed during sexual development (Cordero et al. 1998; Svensson et al. 2009; Willink et al. 2019; Fig. 1).

Previous studies using breeding experiments in controlled laboratory environments across multiple generations have revealed that colour morph development is governed by a single autosomal locus (or a set of tightly linked loci) with sex-limited expression to females (Cordero 1990; Sánchez-Guillén et al. 2005). In *I. elegans*, these female morphs are typically referred to as ‘Androchrome’, ‘Infuscans’ and ‘Infuscans-obsoleta’. To be consistent with previous publications (Willink and Svensson 2017; Willink et al. 2019), we hereafter refer to these three heritable female morphs as A-, I- and O- females.

Shortly after emergence from their last nymphal moult, A- and I-females exhibit a thoracic violet background colour with black antehumeral stripes (Fig. 1b). In A-females, the violet colour becomes turquoise-blue over post-emergence development, and therefore the final thoracic colour pattern of sexually mature A-females strikingly resembles the colour of males (Fig. 1). This adaptive colour change in A-females is consistent with male mimicry, by which A-females avoid excessive and costly male-mating harassment (Gosden and Svensson 2009; Gering 2017). In contrast, the violet thorax colour of I-females becomes greenish-brown in association with increased reproductive output (Willink et al. 2019; Fig. 1b). Finally, O-females are also distinct from males when sexually mature, but they do not exhibit antehumeral stripes at any point in development and their thoracic colour changes from salmon-pink to brown (Cordero et al. 1998; Willink et al. 2019; Fig. 1b).

The two non-mimicking morphs, I- and O-females, exhibit dramatic ontogenetic changes in the phenotypic expression of a blue patch on the eighth abdominal segment (Fig. 1b). This blue abdominal patch is initially present in sexually immature individuals of all three female morphs (Fig. 1b). However, in I- and O- females the blue patch subsequently becomes concealed by dark pigment over sexual development, whereas in males and the male-mimicking A-females, the blue colouration is instead retained during the entire course of sexual development (Henze et al. 2019; Fig. 1). All three female morphs therefore undergo ontogenetic colour changes over sexual development, but there are pronounced differences between A-females and the two other female morphs (I- and O-females). These colour changes occur continuously over the span of a few days (Svensson et al. 2020; Fig. 1b; S1). During this transitional period, individual females with intermediate phenotypes between the immature and mature developmental phases can be visually distinguished (Fig. 1b).

### Fieldwork and sampling design

We sampled three females of each heritable colour morph at three stages of colour development, using the detailed description and classification of these colour stages by Cordero et al. (1998) and Svensson et al. (2009). The *immature* developmental phase was defined in all morphs by the presence of a contrasting blue colour patch in the distal portion of the abdomen, and for A-females by the simultaneous expression of the violet thoracic colouration. Females were classified to the *mature* developmental phase if they expressed the final colour phenotype of sexually mature individuals of their respective morph. Finally, *transitional* females expressed an intermediate phenotype between these two extremes. For I- and O- females these *transitional* individuals were identified by partial concealment of the blue patch with pigment, which results in a brownish-blue abdominal patch (Fig. 1b). *Transitional* A-females were characterised by a violet-turquoise thoracic colouration, which precedes the development of the final turquoise-blue background colour (Fig. 1b). All individuals in this study were collected in the field in southern Sweden during the summer (June 12th – July 7th) of 2015. All individuals were collected during morning hours and from populations that form part of a long-term longitudinal study, in an area of approximately 40 × 40 Km2 (TableS1; Svensson et al. 2005; Le Rouzic et al. 2015; Willink and Svensson 2017).

### RNA sample preparation and sequencing

In addition to the 27 females sampled for the differential gene expression (DGE) analysis, 58 *mature*-coloured females were sampled and used for the *de novo* transcriptome assembly below (Table S1). Field-caught individuals were bisected at the second abdominal segment and the two sections were preserved in RNAlater immediately following bisection, and stored at −80 C within 1-4 h of capture. Samples were then shipped to the Beijing Genomics Institute (1/F, 16th Dai Fu Street, Tai Po Industrial Estate, Tai Po, Hong Kong) for extraction and sequencing. Total RNA was extracted from whole bodies using the Trizol LS Reagent (Life Technologies) following the manufacturer’s instructions. RNAseq libraries were prepared for individual samples using the Illumina TruSeq kit with mRNA enrichment. Each sample was sequenced using 100 bp paired-end reads at a depth of 2 GB per sample on an Illumina HiSeq 2000. Raw data will be deposited at the National Center for Biotechnology Information (NBCI). Sample information and reads will be available through Biosample links and sequence read archives.

### Transcriptome assembly and annotation

Raw reads were trimmed using Trimmomatic v.036 (Bolger et al. 2014) to remove adapter sequences and ambiguous nucleotides. We cropped the first 10 nucleotides of each read and trimmed regions of four nucleotides that had an average quality score below 20. Reads shorter than 25 nucleotides after trimming were discarded. The number of reads per sample after trimming was 9 982 724 ± 1 052 611 (mean ± SD), for all samples used in the DGE analysis. Trimmed reads from all 85 samples were pooled to assemble a reference transcriptome using Trinity v. 2.2.0 (Grabherr et al. 2011). This is the recommended approach for RNAseq analyses on novel species, in order to include more of the alternative splice isoforms in the final reference for mapping (Haas et al. 2013).

Assembly quality was assessed by generating Ex50 estimates using scripts from the Trinity package (Haas et al. 2013), and by quantifying the ortholog hit ratio (OHR) (Vera et al. 2008; O’Neil et al. 2010; O’Neil and Emrich 2013). Ex50 estimates were generated by plotting the N50 of contigs for different expression quantiles, searching for the maximal N50 for regions capturing 70 – 90% of the expression data. OHR estimates were generated using the predicted protein set from the genome of the banded demoiselle damselfly, *Calopteryx splendens* (Csple_OGS_v1.0.faa; Ioannidis et al. 2017), then using blast (tblastn; Camacho et al. 2009) to search for the best hit of each protein in the *I. elegans* transcriptome assembly. OHR was then estimated by dividing the aligned length of an *I. elegans* transcript contig by the length of its orthologous *C. splendens* protein, wherein ratios of 1 indicate the full coverage of the contig for a given protein. The result was a single OHR estimate for each non-redundant protein from a related damselfly, the ortholog of which was found in the *I. elegans* transcriptome. The OHR distribution provides a detailed assessment of a transcriptome, indicating how well thousands of genes have been assembled. Prior to analysis, redundancy due to isoforms and duplications in the *C. splendens* protein set was controlled by collapsing it by the longest protein isoform of each protein cluster group. This was based upon >90% amino acid identity using CD-HIT v4.5.4 (Li and Godzik 2006) with parameters used in making the UniRef90 database (Suzek et al. 2014).

Annotation of the reference transcriptome was conducted using *blastx* v. 2.6.0+ with an e-value of 10e-5 against the National Center for Biotechnology Information (NCBI) non-redundant database (nr). Blast hits were then mapped to Gene Ontology (GO) terms using Blast2GO v. 4.1.9 (Conesa et al. 2005). If the blasted sequences returned more than a single hit, only the highest scoring hit was used in subsequent gene ontology (GO) enrichment analysis.

### Data pre-processing

Prior to the DGE analysis we used the CORSET algorithm (Davidson and Oshlack 2014) to reduce the number of spurious isoforms, which often arise during *de novo* transcriptome assemblies of non-model organisms due to sequencing artefacts. CORSET hierarchically clusters contigs based on shared reads and expression levels. In this case, clusters were based on the 27 samples to be used in our DGE analysis. The read counts are then summarised by these clusters, providing gene-level transcript abundances. Lowly expressed genes (i.e. with fewer than one read per million reads in at least three samples) were discarded at this stage and gene expression distributions were normalised via the weighted trimmed mean of M-values (TMM) method (Robinson and Oshlack 2010).

### Differential gene expression analysis

DGE analyses were conducted using the packages *edgeR* (Robinson et al. 2010; McCarthy et al. 2012) and *limma* (Ritchie et al. 2015) in R v. 3.4.4 (R Core Team 2018), and were based on the workflow described by Law et al. (2016). In order to apply linear-based statistical modelling to the expression data, we estimated the mean-variance relationship in the log transcript counts per million (log-cpm), and incorporated this mean-variance trend into a precision weight for each normalised observation, using the voom method (Law et al. 2014; Liu et al. 2015). The advantage of this statistical approach, which estimates the mean-variance relationship instead of specifying a generating probabilistic distribution, is that it more appropriately controls for type I error rates when sample sizes are small and when sequencing depth varies between samples (Law et al. 2014; Liu et al. 2015). In the *limma* pipeline, these precision weights are then taken into account when fitting the linear models to the log-cpm gene expression data (Law et al. 2016). Empirical Bayes moderation was subsequently applied to these fitted models to more accurately estimate expression variability among genes and the ‘robust’ procedure was employed to allow variance outliers (Smyth 2004; Law et al. 2014; Phipson et al. 2016). Multiple testing across statistical contrasts was conducted using the “global” method, which is recommended when the number of differentially expressed (DE) genes is interpreted as a measure of the strength of a contrast (Smyth et al. 2018). The “global” method performs multiple testing adjustment across the entire set of contrasts and genes being tested. The P-value cut-off was adjusted according to Benjamini and Hochberg’s false discovery rate (Benjamini and Hochberg 1995).

Our goal was to compare DGE across developmental stages and between the different colour morphs. Thus, we fitted a fully-factorial linear model including the fixed effects of female colour morphs, developmental colour phases and their interaction (∼0 + Morph + Phase + Morph:Phase). With this model structure, the intercept is removed from the first factor (Morph), but kept in the second factor (Phase). Therefore, within each morph the immature phase is the intercept, and expression differences between groups can be compared by specifying a contrast matrix. Three types of comparisons (hereafter contrast groups) are possible with the interaction term: (1) contrasts among morphs within each colour development phase, (2) contrasts between subsequent phases within morphs and (3) contrasts among colour morphs in regards to the direction and magnitude of gene expression changes across development (full contrast matrix in Table S2). Because we sampled three females of each of three morphs and at each of three phases in colour development (27 females in total), developmental comparisons in contrast groups (2) and (3) span one of two windows of colour development: *early* colour development, between the *immature* and *transitional* phases, and *late* colour development, between the *transitional* and *mature* phases (Fig. 1b). Our main focus in this study was on contrast group (3), because these contrasts represent changes in expression between developmental colour phases that are opposite in direction or different in magnitude between morphs. We used contrast group (2) to identify genes which are differentially expressed during the course of colour development within morphs. We also used contrasts in (2) to identify the biological processes that are enriched during colour transitions for each female morph. These analyses are summarised in the next two sections. The results of contrasts among morphs within each phase (1) are presented in the Supporting Material (Fig. S2).

The ontogenetic colour changes are not of equal duration in all three morphs (Svensson et al. 2020). I- and O- females take a longer time to complete their colour maturation, compared to A-females (Fig. 1b; S1). As mentioned above, we determined the *transitional* colour phase as being visually intermediate between the *immature* and *mature* colour phases. The *transitional* colour phase is reached approximately at the midpoint of the colour development period in A-females (Fig. 1b). In contrast, I- and O-females that appear intermediate in colour between the *immature* and *mature* colour phases are both older on average than *transitional* A-females, and past the midpoint of their colour development period (Fig. 1b). Consequently, I- and O-females have a protracted *early* colour development compared to A-females, whereas *late* colour development is of a similar duration for all female morphs (Fig. 1b; S1).

### Correlations among subsets of differentially expressed genes

We used vector correlations to determine if developmental expression changes shared similar patterns across morphs for each colour transition (*early* and *late* colour development). Vector correlations are useful to assess the similarity between estimated contrasts of gene expression across a set of multiple genes (Zinna et al. 2018). The correlation between two mean-centred vectors, each containing the estimated magnitude of expression differences between two experimental categories, is:

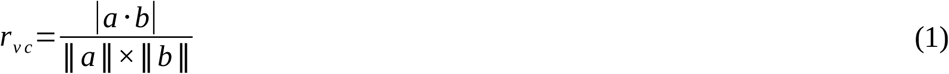

where *a*·*b* is the dot product between the two vectors of statistical contrasts and ‖*a*‖ and ‖*b*‖ represent the magnitudes of vectors *a* and *b* respectively, and are given by the square root of the sum of squares of all contrast estimates. The gene vectors *a* and *b* in this study correspond to log-transformed coefficients estimated from the linear model and contrast matrix described above (Table S2). The vector correlation *r*_*vc*_ is equivalent to the Pearson correlation coefficient, such that values close to 1 indicate an almost perfect correlation between gene expression vectors, values close to 0 indicate no correlation and values close to −1 indicate nearly opposite regulation of gene expression between the two vectors.

We followed the approach of Zinna et al. (2018) to determine if vector correlations between contrasts that have been considered significantly different from zero were particularly strong, compared to correlations between vectors of a random sample of genes, irrespective of the degree of statistical significance. This would indicate that those genes with significant changes in expression across a period of colour development (contrast group (2); Table S2) are also more similar (or dissimilar) in the direction of their regulation than a random gene vector of equal length. We defined in each case an experimental vector, for example, a vector containing all differentially expressed genes during *early* colour development that are shared between two female morphs. We then obtained an empirical distribution of vector correlations, drawn from 10 000 randomly sampled vectors of the same length as the experimental vector. We considered a correlation for the experimental vector as extreme when it fell outside the 95 percentile of correlations in equal-length random vectors.

### Gene ontology enrichment analysis

To uncover the physiological changes that follow colour transitions in the different female morphs, we tested for significantly enriched GO terms in groups of developmentally regulated genes (i.e. genes with significant changes in expression between consecutive colour phases) in comparison to the reference transcriptome, using the bioconductor package TopGO (Alexa and Rahnenfuhrer 2016) in R. For these analyses, we focused on contrasts between developmental colour phases within morphs (contrast group (2); Table S2). In the cases where multiple transcripts associated with a single gene (i.e. a CORSET cluster) differed by at least one GO term (2 231 out of 30 134 mapped transcripts), all GO terms mapped to the different transcripts were pooled, and every unique term was assigned to the gene. Given that our interest was to explore how physiological processes change over development in the three female morphs, here we focused on biological process ontologies. The significance of enriched terms was tested using Fisher’s exact test and the ‘*elim’* algorithm, in order to account for the hierarchical structure of gene ontologies (Alexa et al. 2006). With this algorithm, more general annotations are excluded for genes already counted as significantly enriched for a more specific GO term. GO terms with P-values < 0.01 were considered significantly enriched in the gene set of interest, compared to the reference transcriptome.

### Colour development, heterochrony and female fecundity

A-females reach their final developmental colour phase at a younger age than I- and O-females (Svensson et al. 2020; Fig. 1b). This suggests that unlike colour changes in I- and O-females, colour changes in A-females might be decoupled from sexual maturation. If so, previous records of lower fecundity of A-females compared to I-females (Willink et al. 2019) may be owed to the average younger age of mature-coloured A-females in the field. To test the idea that colour development could be faster (i.e. heterochronic) than most other developmental changes, including sexual maturation, in A-females, we first compared *early* and *late* expression changes within each female morph, using the vector correlation approach described above. A-females reach their mature colour at a similar absolute age as I- and O- females develop their *transitional* phase (Svensson et al. 2020; Fig. 1b). If most other developmental changes proceed at similar rates in all female morphs, we would expect expression contrasts in *early* and *late* colour development to be more correlated in A-females than in the other two morphs, as in A-females these two developmental periods span a shorter absolute time. In contrast, if the fast colour changes of A-females indicate overall faster development, we would expect *early* and *late* expression contrasts to be as different in A-females as in I- and O-females, despite the shorter duration of *early* colour development in A-females.

Furthermore, if A-females accelerate their colour maturation in relation to sexual maturation, we expect that mature-colour A-females in the field would more often be sexually immature, and exhibit a higher rate of reproductive failure than I- and O-females. Our fecundity data to test this prediction come from our surveys of 18 natural populations in Southern Sweden, which were visited frequently during the mating season (June to August) for 2-17 years, between 2001 and 2017 (Svensson et al. 2005; Le Rouzic et al. 2015; Willink and Svensson 2017). During these visits, mating couples were collected in the field. Mated females were placed in individual plastic containers with moist filter paper for oviposition. After 72 h, eggs were scanned at 1 200 dpi and subsequently counted using ImageJ (Schindelin et al. 2012).

In these surveys, mated females were classified by morph and developmental phase. However, *transitional* females were not classified as such, and before 2015 we did not distinguish between A- and I-females in the *immature* developmental phase (Fig. 1b). Therefore, we modelled the probability of reproductive failure only for the *immature* and *mature* developmental phases, and all data on *immature* A- and I-females come from the last three years of our population survey (2015-2017). We used a mixed-effect model fitted by MCMC in the *R* package MCMCglmm (Hadfield 2010) to test for a difference between morphs in the probability of reproductive failure, which we defined here as mated females that produced no viable eggs. We specified a binomial model with morph, developmental phase and their interaction as fixed effects. We also allowed for a random interaction between the population and season terms to affect the variance around the fixed effect estimates. All code to reproduce these analyses as well as the reproductive failure data will be uploaded to Dryad.

## Results

### Reference transcriptome

Using an extensive collection of RNA-Seq data, we assembled a transcriptome of 889,001 transcripts. Grouping transcripts by their expression levels and looking at those with the highest levels of expression in relation to their assembly lengths, a peak of transcript assembly length (N50) of nearly 3kb was found at expression levels containing 70-80% of transcripts (i.e. ExN50 ∼ 3kb), suggesting sufficient data for a high quality assembly. We also estimated the OHR of our assembly by querying our assembly with the full protein set of the only published odonate genome, the banded demoiselle damselfly (*Calopteryx splendens*), which shared a common ancestor with *I. elegans* approximately 85 million years ago (the average from two dated phylogenies, 92 Ma in Thomas et al. 2013 and 79 million years in Waller and Svensson 2017). Of the 22,507 *C. splendens* proteins that were non-redundant, we found good hits (e-value less than 10e-5) for roughly 19,962, and of these, we found OHR of 0.7 and higher for nearly 70% (13,497; see Fig. S2 and Table S3). Given the extensive divergence between the two taxa, the number of putative orthologs between species recovered is high and the strong skew towards high OHR values is indicative of a high quality transcriptome assembly.

### General trends in early colour development

In all female morphs, *early* colour development was characterised by an extensive decrease in gene expression between the *immature* and *transitional* phases (contrast group (2); Table 1; Table S2). Across-the-board downregulation from the *immature* to the *transitional* phase exceeded upregulation by a factor of more than 4, in O-females, to more than 25, in A-females (Table 1). Of all downregulated genes during *early* colour development, 3 158 were shared by the three female morphs, whereas there were no upregulated genes shared by all morphs. The downregulated genes shared by all female morphs corresponded to ∼ 62%, 75% and 67% of all downregulated genes in A-, I- and O- females, respectively. These genes were significantly enriched for biological processes involved in protein synthesis, transport and catabolism, energy metabolism, signal transduction, and cathecolamine biosynthesis (Table S4). Broadly shared downregulation of gene expression across *early* colour development also resulted in highly correlated vectors of genes with significant expression changes between all morph pairs (Fig. 2a).

**Table 1.** Number of differentially expressed (upregulated and downregulated) and similarly expressed (N.S.) genes, in comparisons between developmental colour phases and within female colour morphs of *I. elegans*. Developmental comparisons are performed between the *immature* and *transitional* phases (*early* colour development) and between the *transitional* and *mature* developmental phases (*late* colour development). A total of 27 females were sampled for this analysis, three of each colour morph in all three developmental phases.

| Colour transition | Morph | Upregulated | Downregulated | N.S. |
| --- | --- | --- | --- | --- |
| Early | A | 197 | 5 200 | 20 983 |
|  | I | 302 | 4 190 | 21 888 |
|  | O | 978 | 4 685 | 20 717 |
| Late | A | 3 256 | 1 731 | 21 393 |
|  | I | 364 | 355 | 25 661 |
|  | O | 239 | 471 | 25 670 |

**Figure 2.**
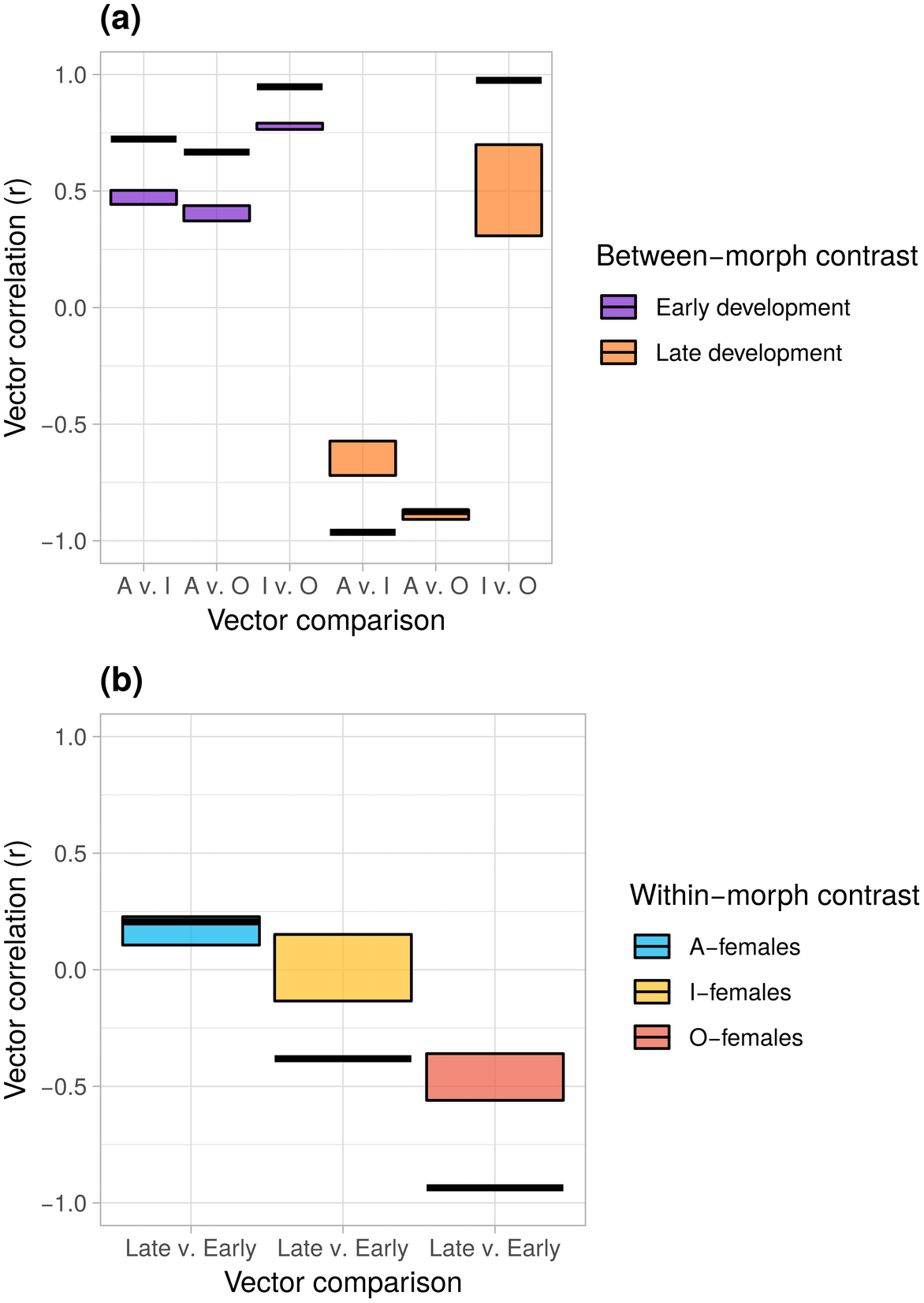
Vector correlations of gene expression contrasts. We tested for overall similarity in the expression changes underlying colour development **(a)** between all female-morph pairwise comparisons during *early* and *late* colour development, and **(b)** between *early* and *late* colour development within each female morph. Vector correlations (r) indicate the similarity in the direction of change in gene expression between two vectors of developmental contrasts. Values close to 1 suggest that the two vectors are nearly identical in direction, whereas values close to −1 suggest developmental regulation of gene expression is opposite in direction between the two vectors. The black horizontal lines show correlation values for experimental vectors (sets of differentially expressed genes, see Methods), the coloured boxes show the 95 percentile of empirical distributions of 10 000 contrast vectors of the same length as the experimental vectors, but including a random sample of all genes. The lengths of the experimental gene vector from left to right are 3528, 3589, 3785, 269, 652 and 66 for **(a)** and 1051, 186, 221 for **(b)**.

### General trends in late colour development

No significant expression changes from the *transitional* to the *mature* colour phase (contrast group (2); Table S2) were shared by all female morphs. This absence of shared regulatory patterns during *late* colour development was caused by the distinctiveness of A-females. There was a perfect negative correlation between A- and I-females in the regulation of genes that were differentially expressed in both morphs (Fig. 2a). Of 269 genes with significant expression changes between the *transitional* and *mature* colour phases, 164 increased in expression in I-females and decreased in expression in A-females, whereas 105 exhibited the opposite regulatory pattern. There was also a negative correlation in the direction of regulation for the genes differentially expressed across *late* colour development in A- and O-females, but this correlation was not particularly extreme (P = 0.14). Most genes, whether significant or not, exhibited opposite developmental contrasts between A- and O-females during *late* colour development. In contrast, there were 66 genes with significant changes in expression across *late* colour development in both I and O-females, and all of these genes were regulated in the same direction in both morphs (i.e. either increased or decreased in expression between the *transitional* and *mature* colour phases; Fig. 2a).

The biological processes enriched in DE genes during *late* colour development were also markedly different between A-females and both I- and O-females. Compared to the *transitional* colour phase, *mature*-colour A-females upregulated genes enriched for functions in cell cycle progression, meiosis and the regulation of transcription, while downregulating multiple processes associated with energy metabolism (Table S6; S7). Both I- and O-females in their *mature* colour phase upregulated energy metabolic processes compared to their *transitional* colour phase (Table S7). *Mature*-colour I-females downregulated lipid metabolism and gene silencing and *mature*-colour O-females downregulated genes participating in cell cycle progression, protein and RNA modification, compared to females of these morphs in the *transitional* colour phase (Table S6).

### Colour development heterochrony and female fecundity

I- and O-females underwent a qualitative change in the direction of gene regulation between *early* and *late* colour development. DE genes that increased in expression between the *immature* and *transitional* colour phase (*early* colour development) tended to decrease in expression between the *transitional* and *mature* colour phase (*late* colour development), and *vice versa* (Fig. 2b). In contrast, regulation of gene expression was positively, although not extremely, correlated between *early* and *late* colour development in A-females (Fig. 2b). Such positive correlation between developmental periods in A-females was not extreme when compared to a random sample of expression contrasts drawn from equal-length gene vectors regardless of their significance level (P = 0.163; Fig. 2b). Nonetheless, DE genes that decreased in expression through both *early* and *late* colour development in A-females were enriched for two shared biological processes, both associated with ATP biosynthesis (Table S4; S6). DE genes across *early* and *late* colour development did not share any enriched biological processes for either I- or O-females.

We obtained data on reproductive failure of 5 247 (NA = 3 336, NI = 1 667, NO = 234) *mature*-colour and 93 (NA = 28, NI = 34, NO = 21) *immature*-colour females, all mated in the field. Across these three female morphs, *immature* developmental phases exhibited a nearly four-fold probability of reproductive failure compared to their *mature*-colour counterparts (PMCMC < 0.001; Fig. 3). There were no differences in the rate of reproductive failure among the three female morphs in the immature phase (all PMCMC > 0.05; Fig. 3a). Among mature-colour individuals, reproductive failure was somewhat higher in A-females compared to I-females (PMCMC = 0.019; Fig. 3b), yet this difference was small relative to differences between mature and immature-coloured females (Fig. 3).

**Figure 3.**
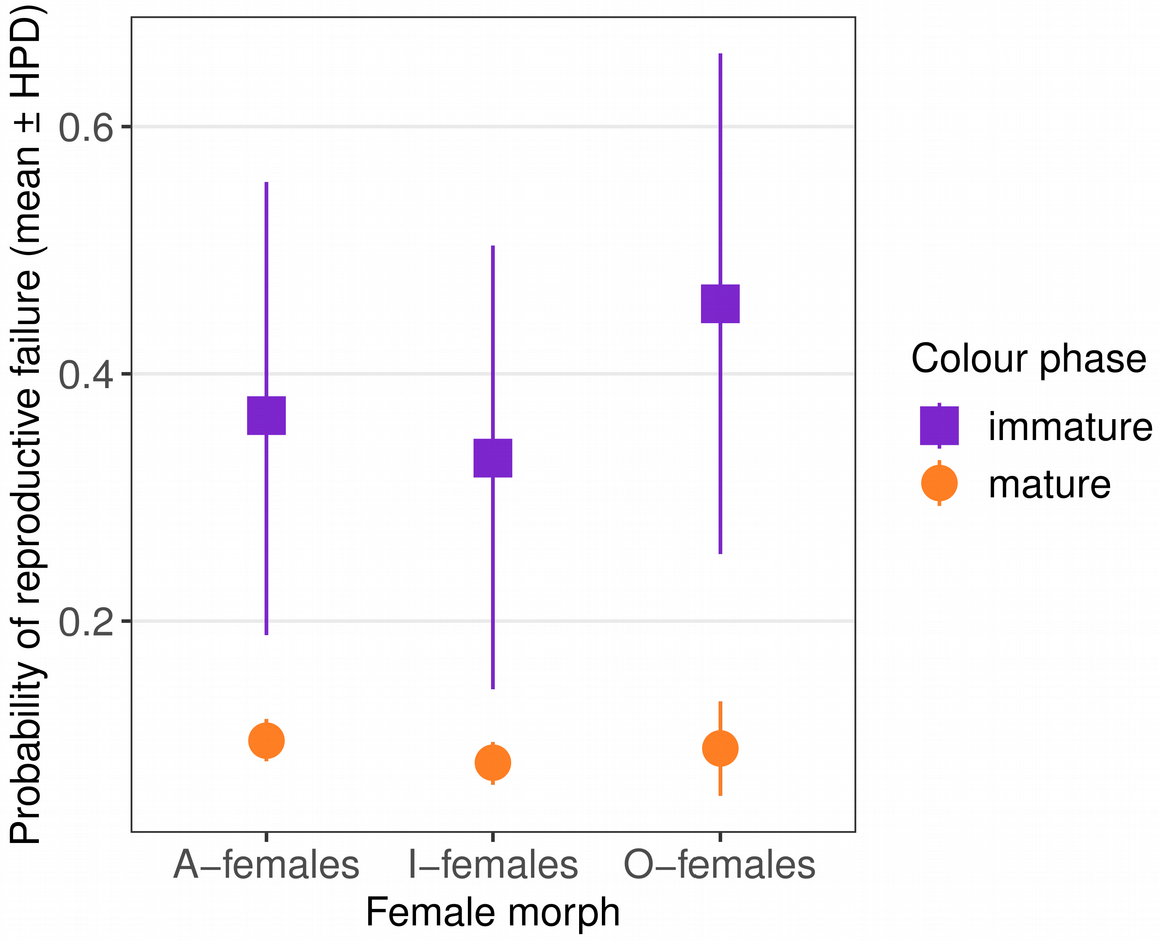
Probability of reproductive failure, quantified as the proportion of mated females laying no viable eggs, among three female morphs and two developmental colour phases of *I. elegans*. The symbols show the posterior mean and 95% highest posterior density (HPD) intervals of parameter estimates in a binomial mixed-effect model fitted by MCMC.

### Regulatory differentiation during colour development

We tested whether changes in gene expression between consecutive developmental colour phases (i.e. developmental regulation) differed in direction or magnitude between female morphs (contrast group (3); Table S2). There were more substantial morph differences in developmental regulation of gene expression during the second phase of colour development (*late* colour development; Fig. 4). During this period, A-females were differentiated from both I- and O-females in the pattern of developmental regulation of 856 genes (Fig. 4). Statistical contrasts for all these genes were of the same direction between A- and I-females and between A- and O-females.

**Figure 4.**
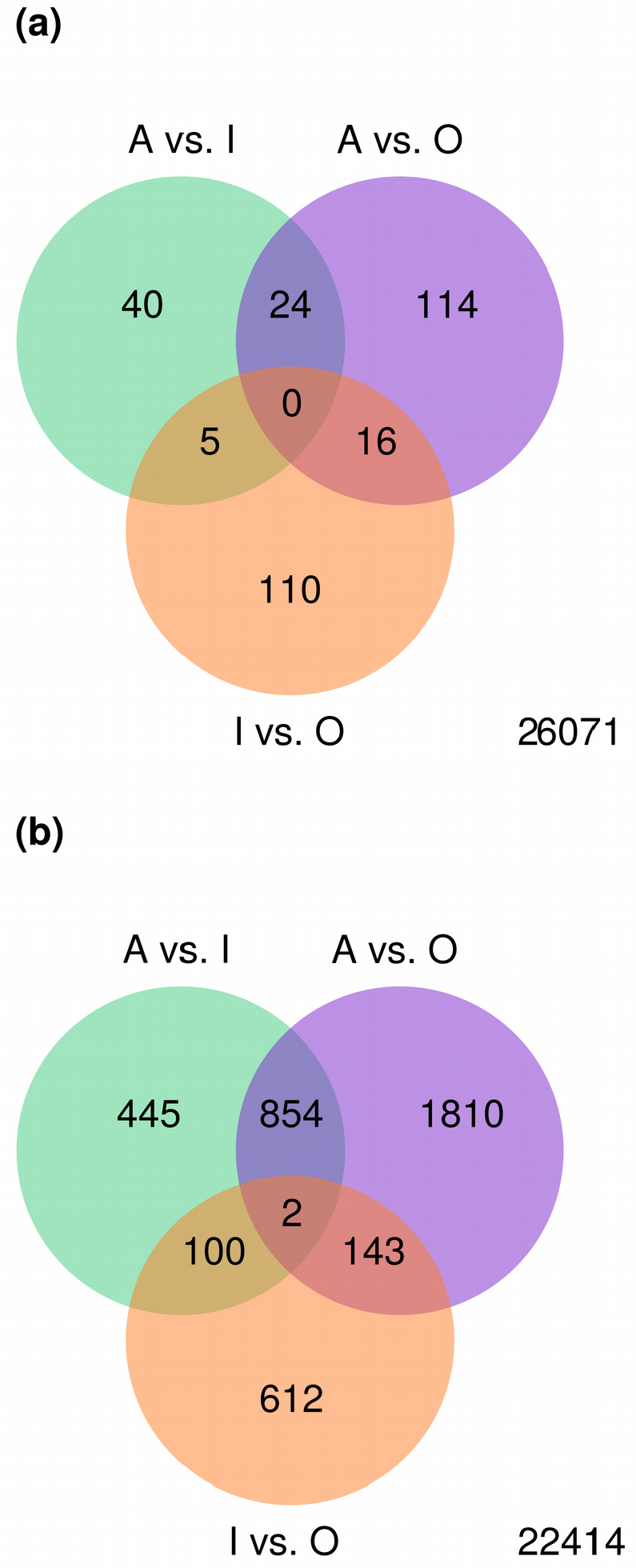
Venn diagram showing the genes that were differentially regulated across colour development, between pairs of female morphs of *I. elegans* (i.e. genes with significant differences between morphs in the direction or magnitude of developmental changes; contrast group (3); Table S2). **(a)** *early* colour development, **(b)** *late* colour development. The number of genes with no evidence of morph differences in differential expression between developmental colour phases is shown in the bottom right corner of each plot.

To examine how epistatic effects of the colour-morph locus could mediate the differentiation of A-females, we queried subsets of developmentally regulated genes for regulatory functions. Here, we focused on genes for which the direction or magnitude of developmental regulation differed significantly between A- and both I- and O-females (contrast group (3); Table S2). First, we used the search term ‘juvenile hormone’ (JH), and ‘ecdysone’ (E), two major hormones involved in regulating insect reproduction and development (Simonet et al. 2004; Flatt et al. 2005). While we found no differentially regulated genes directly involved in JH metabolism, four genes related to ecdysone metabolism and signalling exhibited a different developmental change in expression in A-females compared to I- and O-females, during *late* colour development. None of these genes, or any other ecdysone-related gene, differed in developmental regulation between I- and O- females during *late* colour development, and neither did they differ between any pair of morphs during *early* colour development. To discern the potential functions of these genes, we used *blastx* against annotated proteins in the *Drosophila melanogaster* genome in FlyBase (Thurmond et al. 2019). Three of these ecdysone-related genes were annotated to *D. melanogaster* genes in the *Halloween* group, which code for Cytochrome P450 enzymes that participate in ecdysone biosynthesis (Table 2; Fig. 5). The fourth gene was annotated as coding for the ecdysone-induced protein E74, a transcription factor that responds to concentrations of 20-hydroxyecdysone (20E) in the ovary of *D. melanogaster* (Table 3; Fig. 6; Ables and Drummond-Barbosa 2010).

**Table 2.**
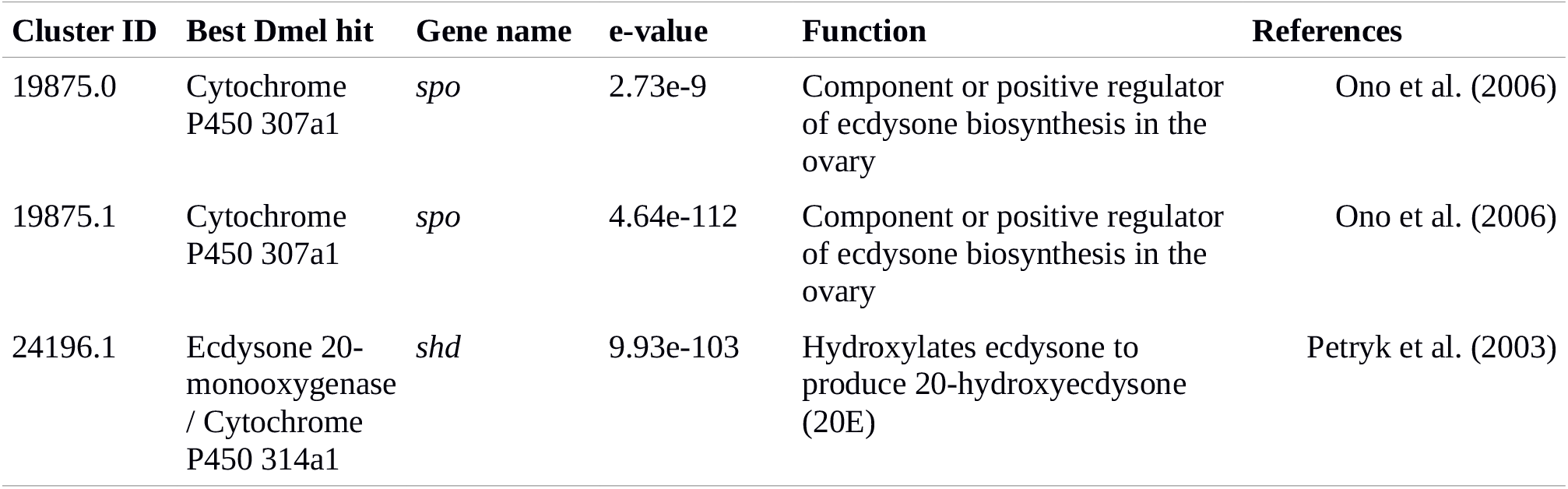
Ecdysone metabolism genes with different changes in gene expression between A- and both I- and O-female morphs in *I. elegans*, during *late* colour development. CORSET clusters are assumed to represent *I. elegans* genes. Transcript sequences in these clusters were blasted against annotated proteins in the *Drosophila melanogaster* (Dmel) genome. For each of the queried *I. elegans* sequences, we report the best Dmel hit, its corresponding gene name and inferred function.

**Table 3.** Regulatory genes with different changes in gene expression between A- and both I- and O-female morphs in *I. elegans*, during *late* colour development. CORSET clusters are assumed to represent *I. elegans* genes. Transcript sequences in these clusters were blasted against annotated proteins in the *Drosophila melanogaster* (Dmel) genome. For each of the queried *I. elegans* sequences, we report the best Dmel hit, its corresponding gene name, e-value and inferred mode of regulation and function.

| Cluster ID | Best Dmel hit | Gene name | e-value | Mode of regulation | Function | References |
| --- | --- | --- | --- | --- | --- | --- |
| 16360.1 | Transcription termination factor, mitochondrial | *mTTF* | 4.81e-32 | Transcription factor | Regulates mitochondrial replication and transcription by interacting physically with helicase and polymerase enzymes | Roberti et al. (2005) |
| 18749.0 | T-related protein | *brachyenteron* | 2.09e-49 | Transcription factor | Fate determination of hindgut stem cells | Fox and Spradling (2009) |
| 19883.1 | Histone deacetylase complex subunit SAP30 homolog | *Sap30* | 2.71e-58 | Post-translational modification | Part of a corepressor complex that regulates G2/M cell cycle progression | Laherty et al. (1998) |
| 28078.0 | Ecdysone-induced protein E74 | *Eip74EF* | 1.01e-52 | Transcription factor | Promotes germline proliferation and maintenance in ovaries | Ables and Drummond-Barbosa (2010) |
| 31494.0 | Enhancer of split mbeta protein | *E(spl)mbeta-HLH* | 7.48e-20 | Transcription factor | Repression of neural cell fate in response to Notch signalling | Jennings et al. (1994) |
| 36913.0 | Vrille | *vri* | 7.43e-46 | Transcription factor | Control of circadian rhythm, cardiac aging, ecdysone target in the ovary, associated with follicle cell maintenance and proliferation | Cyran et al. (2003); Monnier et al. (2012); Ables et al. (2016) |
| 41360.2 | Chromodomain-helicase-DNA-binding protein Mi-2 homolog | *Mi-2* | ~ 0 | Chromatin remodelling | Proneural gene repression | Yamasaki and Nishida (2006) |
| 42782.0 | Bunched | *bun* | 6.00e-31 | Transcription factor | Ovarian follicle cell development and migration, eggshell formation, neurblast proliferation, mushroom body development | Dobens et al. (1997); Dobens et al. (2000); Kim et al. (2009) |
| 44536.0 | Traffic jam | *tj* | 2.66e-29 | Transcription factor | Ovarian follicle cell migration, maintenance of ovarian stem cells | Gunawan et al. (2013); Panchal et al. (2017) |
| 44568.1 | Histone acetyltransferase | *enok* | 8.66e-110 | Post-translational modification | Neuroblast proliferation, maintenance of ovarian germ cells | Scott et al. (2001); Xin et al. (2013) |
| 46100.0 | Heterogeneous nuclear ribonucleoprotein A1 | *Hrb98DE* | 2.93e-36 | mRNA splicing and export | maintenance and differentiation of ovarian germ cells, | He and Smith (2009); Ji and Tulin (2012) |
| 46510.1 | Homeotic protein Sex combs reduced | *Scr* | 3.80e-46 | Transcription factor | Induces and responds to the expression of *dsx* during sexual differentiation of somatic tissues | Barmina and Kopp (2007); Tanaka et al. (2011) |
| 47599.1 | Doublesex-Mab related 99B | *dmrt99B* | 5.82e-20 | Transcription factor | Integrates sex-determining, patterning and hormonal signals during somatic sexual differentiation | Kopp (2012) |

**Figure 5.**
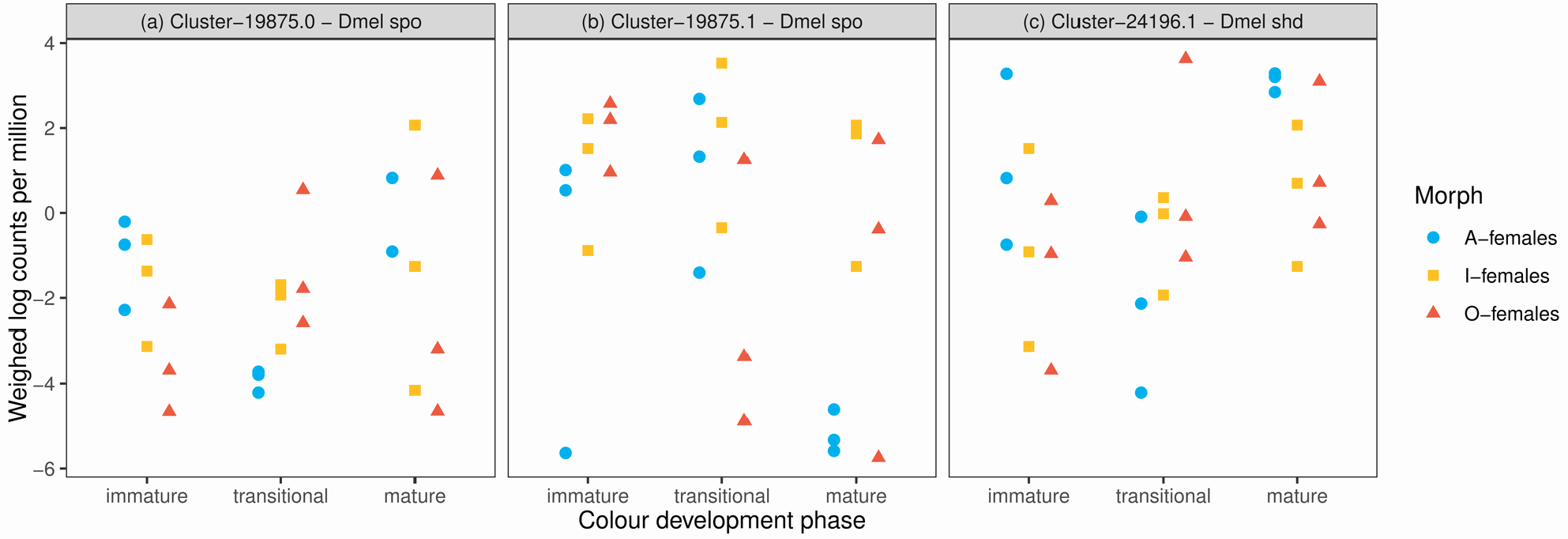
Expression patterns of ecdysone metabolism genes with different changes in gene expression between A- and both I- and O-females during *late* colour development of *I. elegans*. Expression data (log counts per million) have been weighed by within-sample variability and by the mean-variance relationship among genes, using ‘voom’ (see Methods). Best hits in the *D. melanogaster* (Dmel) genome are shown in Table 2.

**Figure 6.**
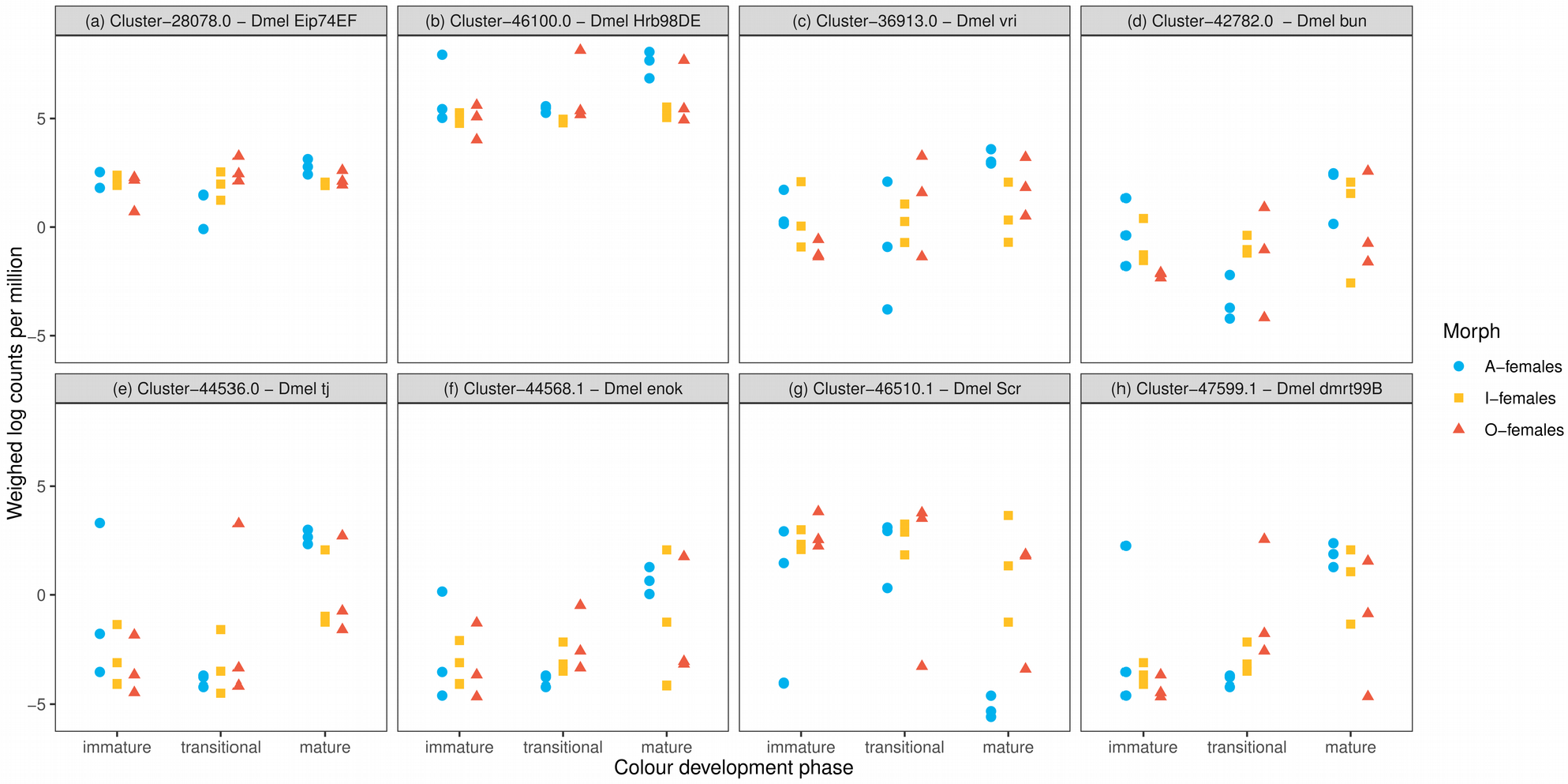
Regulatory genes with different changes in gene expression between A- and both I- and O-females, during *late* colour development of *I. elegans.* The circles indicate expression levels of genes with inferred roles in regulating reproductive development (**a-f**) and sexual differentiation (**g-h**) in *D. melanogaster*. Expression data (log counts per million) have been weighed by within-sample variability and by the mean-variance relationship among genes, using ‘voom’ (see Methods). Best hits in the *D. melanogaster* (Dmel) genome and inferred modes of regulation and functions are shown in Table 3.

We further identified all genes annotated with the GO term “regulation of transcription”, within the gene set that underwent unique developmental changes in expression in A-females during *late* colour development (Fig. 4b). This query revealed 18 genes (Table 3), most of which (∼89%) increased in expression in A-females, while remaining relatively constant or decreasing in I- and O-females. We again used *blastx* against the *D. melanogaster* genome to examine the potential functions of these regulatory genes. We focus on the 13 genes that had good hits (e-value < 10e-5) against annotated proteins in SwissProt/TrEMBL (UniProt Consortium 2019). These genes comprised several transcription factors, including E74, a ribonucleoprotein and two proteins involved in histone acetylation (Table 3; Fig. 6). None of these regulatory genes varied significantly in expression patterns between I- and O-females, during either *early* or *late* colour development.

## Discussion

The genomic signatures and consequences of sexual antagonism and various forms of sexual conflict is gaining increased interest (Rowe et al. 2019), but with a few recent exceptions from studies on fruitflies, humans and flycatchers, empirical data in this area is limited (Barson et al. 2015; Cheng and Kirkpatrick 2016; Dutoit et al. 2018). We suggest that research in this area would benefit from gene expression studies on organisms with discrete genetic morphs, including female-limited polymorphisms, which are many times associated with sexual conflict between males and females (Svensson et al. 2005; 2009; Reinhardt et al. 2007; Karlsson et al. 2013; Iversen et al. 2019). Here, we have presented data on how different female morphs of the damselfly *I. elegans* change their gene expression profiles in different ways during the course of colour development, as these morphs become sexually mature (Willink et al. 2019). One of the female morphs develops a male-like colouration at maturity, whereas the other two morphs develop colour patterns strikingly different from that of males (Fig. 1b; Svensson et al. 2009). Morph differences in the direction, magnitude or timing of gene expression changes during this developmental period could suggest direct or indirect interactions between the colour-morph locus and loci underlying the multitude of phenotypic traits that differentiate the adult female morphs (Svensson and Abbott 2005; Lancaster et al. 2017; Willink and Svensson 2017; Willink et al. 2019; Svensson et al. 2020). Elucidating these interactions is important for a better understanding of the mechanisms behind pleiotropy of major-effect loci, and how these loci can produce phenotypically divergent and co-adapted morphs in damselflies and other insects.

In this study, we found that during the *early* phase of colour development, between the *immature* and *transitional* phases, thousands of genes, controlling multiple metabolic processes, decreased in expression in all three female morphs (Table 1; S4; Fig. 2a). Widespread downregulation of gene expression may indicate a shared slowdown in basic metabolism across all three morphs, which characterises adult development in insects once energetically expensive tissues, such as gametes and flight muscle, have developed (Hack 1997; Piiroinen et al. 2010; Niitepõld and Hanski 2013). Conversely, most differences in gene expression changes among morphs were observed *late* in colour development, between the *transitional* and *mature* developmental colour phases (Fig. 4). During *late* colour development, trajectories of gene expression changes diverged markedly between the male-mimicking A-females and both I- and O-females (Fig. 4b). I- and O-females also exhibited greater differentiation in developmental regulation of gene expression during *late* colour development, but neither morph was as differentiated from the other two as A-females were (Fig. 4b). Moreover, the overall expression patterns of DE genes in I- and O-females remained positively correlated, in *late* as in *early* colour development (Fig. 2a).

The male-mimicking A-females of *I. elegans* develop their final colour pattern at an earlier age than the other female morphs (Svensson et al. 2020; Fig. 1b; S1). Thus, one possible explanation for the distinctive gene expression profiles in A-females is that they accelerate colour changes relative to other developmental processes, including sexual maturation. If so, A-females would become differentiated from I- and O-females during *late* colour development primarily because of their younger age. Our results suggested that colour development in A-females is indeed accelerated with respect to some other developmental processes, but not necessarily sexual maturation. DE genes across *early* and *late* colour transitions were positively correlated in A-females only (Fig. 2b), as expected by the fact that the two colour transitions occur over a shorter time period in this morph. However, this correlation was not particularly extreme, when compared to a random sample of genes (Fig. 2b). *Mature*-colour A-females showed a somewhat increased tendency to exhibit reproductive failure compared to *mature*-colour I-females, as expected if a larger proportion of *mature*-colour A-females was still sexually immature. However, this difference was relatively small (less than 2% on average; Fig. 3), and it is therefore unlikely to account for the large differences in fecundity between A- and I-females in the field (Willink and Svensson 2017; Willink et al. 2019). Taken together with our recent previous work, these results suggest that heterochrony of colour development relative to other developmental processes may explain some of the differences in fecundity and gene expression between A- and I-females. Yet, sexual maturation is also accelerated in A-females. In fact, these male-mimicking females achieved a relatively high rate of reproductive success after only two thirds of the colour-development period of I- and O-females (Fig. 1b; 3). The reduced fecundity of the male-mimicking A-females might therefore be a pleiotropic cost caused by heterochrony, that this morph pays for its accelerated colour development. However, the results in the present study do not permit us to pinpoint the specific physiological or molecular mechanisms behind such a cost. The accelerated reproductive development in A-females might however be advantageous, especially at northern latitudes where mating seasons are cooler and shorter (Svensson et al. 2020).

Some previous studies on vertebrates suggest that a single locus or a tightly linked group of colour-morph loci can influence reproductive physiology via hormonal regulation (Sinervo and Calsbeek 2003; Lamichhaney et al. 2016). In *I. elegans*, the colour-morph locus might also interact via hormonal regulation with numerous other genes that influence egg development. We found that three *I. elegans* genes with inferred roles in the ecdysone biosynthesis pathway underwent changes in expression during *late* colour development that were more pronounced (i.e. either more negative or more positive) in A-females than in the other two female morphs. One of these genes was inferred to code for 20EMO, the enzyme that catalyses the conversion of E to 20E in the ovarian follicle and nurse cells of *D. melanogaster* (Table 2; Fig. 5c; Petryk et al. 2003). In insects, 20E induces ovarian cell development and is required in follicle cell differentiation and vitellogenesis (Swevers and Iatrou 2003; Simonet et al. 2004). In A-females of *I. elegans*, this putative gene increased in expression between the *transitional* and *mature* colour phases, reaching an expression level similar to that of sexually mature I- and O-females, which are older on average (Fig. 1b; 5c). The two other genes with unique developmental changes in expression in A-females code for enzymes associated with upstream points in the ecdysone biosynthesis pathway (Table 2; Fig. 5a, b). Finally, an ecdysone responsive gene essential for oogenesis (E74) also showed increased expression during *late* colour development in only A-females (Table 4; Fig. 6a). In *D. melanogaster*, E74 responds to ecdysone titres to promote germline proliferation during early oogenesis, and is key for preventing germline degeneration during later stages of oogenesis (Ables and Drummond-Barbosa 2010).

Our finding that both ecdysone biosynthesis and ecdysone-responsive genes changed in expression during the *late* colour development of only A-females (Fig. 5; 6a) suggests that the colour morph locus might have pleiotropic effects via ecdysone signalling, regulating the timing of female reproductive development. This interpretation is also supported by the expression pattern of several other regulatory genes that are presumably involved in reproductive development (Table 3), all of which increased in expression between the *transitional* and *mature* colour phases of only A-females (Fig. 6b-f). For instance, in *D. melanogaster*, a heterogeneous ribonucleoprotein (hnRNP) induces the expression of cadherin, which is in turn essential to maintain cell-cell adhesion during germline stem cell proliferation (Ji and Tulin 2012). hnRNPs can be responsive to ecdysone levels through association with the protein on ecdysone puffs (PEP) (Amero et al. 1993). In *I. elegans*, both hnRNP and cadherin genes increased in expression during the *late* colour development of adult A-females (Fig. 6b; S4). In fact, *mature* colour A-females had a higher expression level of this putative hnRNP gene than either I- or O-females (Fig. 6b). Also consistent with the accelerated sexual maturation in A-females, the set of genes that increased in expression during *late* colour development in this morph was significantly enriched for multiple processes necessary for cell cycle progression, including meiotic cell cycle and division (Table S7). This was not the case for I- and O-females during the course of either *early* or *late* colour development.

The regulatory genes that were differentially expressed in the male-mimicking A-females during *late* colour development also suggest a role of the colour-morph locus in inducing morph differentiation in somatic tissues of *I. elegans*. One of these genes mapped to *Dmrt* (Table 3), which is a gene family involved in sex determination and differentiation across animal taxa (Kopp 2012). This *Dmrt* gene underwent a greater increase in expression between the *transitional* and *mature* colour phases in A-females compared to both I- and O-females. In somatic tissues of *D. melanogaster* another *Dmrt* gene, *doublesex* (*dsx*) integrates sexual identity, patterning and hormonal signals to induce the development of sexually dimorphic structures (Kopp 2012). Alternative splicing of *dsx* also underpins the development of male-mimicry in the female polymorphic butterfly *Papilio polytes* (Kunte et al. 2014). In the *I. elegans* congener *I. senegalensis*, non-mimicking females are distinguished from both A-females and males in the expression pattern of a *dsx* isoform (Takahashi et al. 2018). *dsx* is therefore also a plausible regulator of the development of male mimicry in *I. elegans,* as it is differentially expressed between adult males and females (Chauhan et al. unpublished manuscript). However, we found no evidence of differential expression of the *Dmrt* gene *dsx* in *I. elegans* females, during the period of developmental colour changes (Fig. S5a). The *Dmrt* gene differentially expressed during *late* colour development in *I. elegans* lacks homology to the dimerization domain that characterises *dsx* (Bayrer et al. 2005), but as all other *Dmrt* genes it contains a DM DNA binding motif, which confers regulatory capacity over pseudopalindromic DNA targets (Zhu et al. 2000).

Holometabolous insects have multiple *Dmrt* genes (Wexler et al. 2014), and *I. elegans* has at least two (Fig. 6h; S5a). However, the functions of the *Dmrt* genes other than *dsx* are poorly known in insects, although these genes might also integrate sexual identity, spatial and temporal signals during sexual differentiation (Picard et al. 2015). While sex determination mechanisms are diverse across animal taxa, their downstream actions which result in sexual differentiation are typically more conserved and often rely on transcriptional regulation by the DM domain DNA binding motif (Zhu et al. 2000). In line with the possibility that this *Dmrt* gene may contribute to the somatic differentiation of male-mimicking females in *I. elegans*, we found that genes which interact with *dsx* during the development of sexually dimorphic structures in *D. melanogaster* also had more drastic changes in expression during *late* colour development in A-females than in either I- and O-females (Table 4; Fig. 6g; S5b-d).

This study of the developmental trajectories of colour and gene expression changes in the female morphs of *I. elegans* contributes to understanding the mechanisms for pleiotropy of large-effect loci, such as those underpinning animal colour polymorphisms. Our results suggest that the colour-morph locus in *I. elegans* induces heterochrony in the male-mimicking A-females. These A-females show accelerated colour changes and faster sexual maturation, the latter presumably caused by ecdysone signalling. We also uncovered hundreds of developmental changes in gene expression profiles that differed in direction or magnitude between A-females and both I- and O-females (Fig. 4b). There are three possible explanations for this pattern. First, some differences in gene expression changes during *late* colour development may arise if *mature*-colour A-females were younger than the other *mature*-colour females. This could explain phenotypic differences in traits other than colour and fecundity, both of which develop faster in A-females. Second, developmental genes with many downstream targets, such as *Dmrt* and homeotic genes (Table 4; Fig. 6g-h), may mediate unique expression changes of A-females through complex networks of gene-gene interactions. Third, the colour locus may cause ecological differentiation between morphs via fewer direct interactions between genes, but with morph-specific micro-habitat selection, differential parasite loads or differential inter-sexual interactions feeding back into the developmental regulation of gene expression. The latter two possibilities represent functional epistasis, whereby the phenotypic effects of genes or gene substitutions depend on the genetic background (Hansen and Wagner 2001), in this case, on the genotype at the colour-morph locus. Theory shows that functional epistasis can play crucial roles in evolution by natural selection (e.g. Kimura and Maruyama 1966; Barton 1995; Rice 1998; Hansen 2013; Jones et al. 2014). However, in order to gauge its empirical importance we need to better understand the dynamics of gene interactions over the course of development (Csilléry et al. 2018).

## Conclusions

Female morphs of the common bluetail damselfly differ in their developmental trajectories of colour change, sexual maturation and changes in gene expression. Specifically, the male-mimicking A-female morph exhibits accelerated colour and reproductive development, and becomes increasingly different from the two other female morphs in the pattern of developmental changes in gene expression, soon before the completion of colour development. Our results suggest that ecdysone signalling mediates epistatic interactions between the colour-morph locus and genes influencing the development of female fecundity. Our results also point to other regulatory differences during this critical period of colour development, such as a drastically increased expression of a *Dmrt* gene during the development of male-mimicking females. This study provides insights on how pronounced phenotypic differences between female morphs emerge during the course of development, and suggests that heterochrony and epistasis may underlie such pleiotropic effects of colour morph loci in *I. elegans*. These results add to a growing understanding of how the evolution of regulatory architecture is shaped by sexual conflict.

## Acknowledgements

Funding for this study have been provided by research grants from The Swedish Research Council (VR: grant no. 2016-03356) and Erik Philip Sörensens Stiftelse to E.I.S. M.C.D was supported by an NSF Postdoctoral Research Fellowship in Biology (DBI 1401862). We are also grateful to the numerous field assistants and students, who participated in the field surveys of damselfly populations over the last two decades, and to the Swedish National Infrastructure for Computing (SNIC) for providing us with computational resources. We thank Rachel Blow, Stephen De Lisle, Miguel Gómez-Llano and two anonymous reviewers for helpful comments on earlier versions of this manuscript.

## Author contributions

E.I.S. conceived the study with input from M.C.D. and B.W. M.C.D. and B.W. obtained the sequence data. C.W. assembled the trancriptome. B.W. performed the differential expression analyses and functional annotations with help from M.C.D. B.W. wrote the manuscript with input from all authors.

