## Supporting Material for "Changes in gene expression during female reproductive development in a colour polymorphic insect"

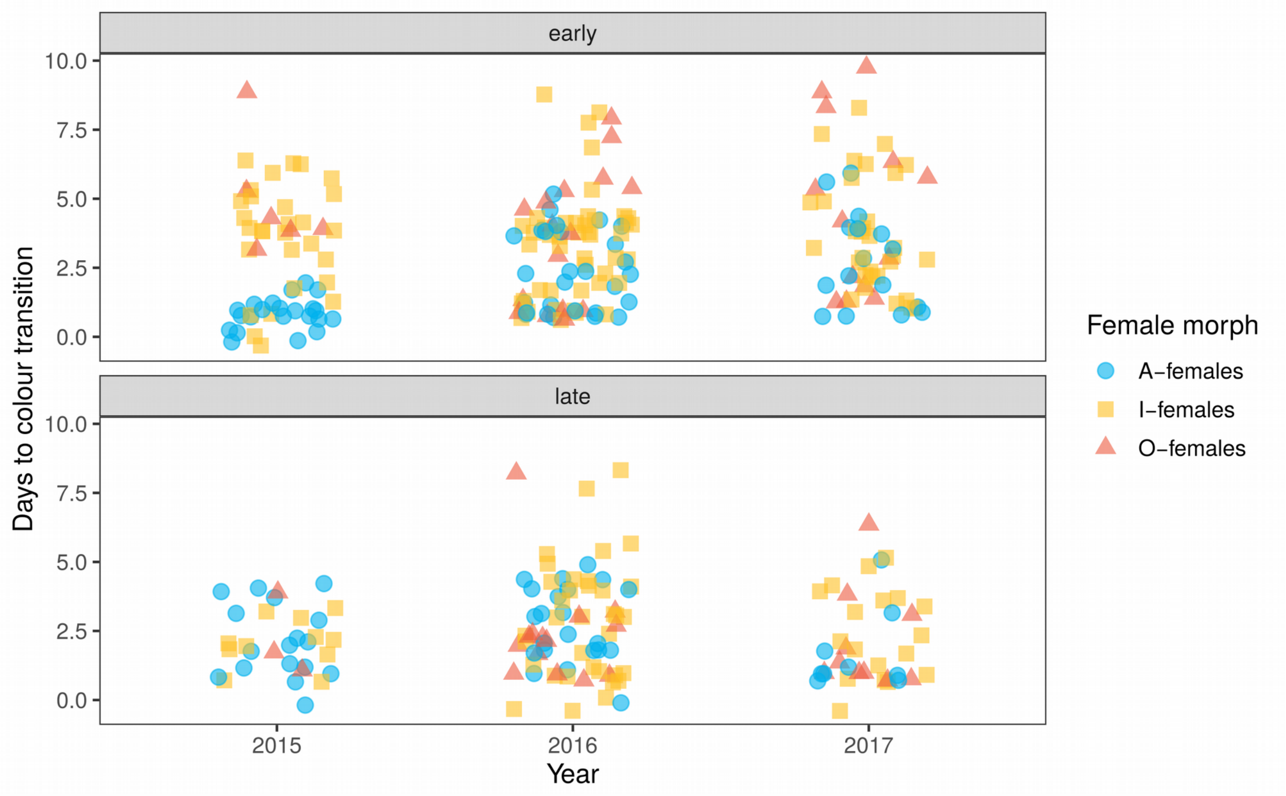


**Figure S1.** Duration of colour development in the three female morphs of *Ischnura elegans*. A total of 212 emerged females ($N_A = 66$ , $N_I = 108$, $N_O = 38$) were raised in outdoor enclosures and the first day of observation of each developmental colour phase was recorded for surviving individuals. The points show the number of days to reach the *transitional* colour phase (end of *early* colour development) and the *mature* colour phase (end of *late* colour development), for females raised on three consecutive years. A mixed effect model fitted by MCMC, with time-to-colour-phase as the response variable and the interaction between morph (A- I- or O-females), colour development (*early* or *late*) and year (2015, 2016 or 2017) as predictor variables, uncovered a significantly shorter duration of *early* colour development of A-females compared to I- and O-females in 2015 ($PMCM_{A\ vs\ I} < 0.001$, $PMCMC_{A\ vs\ O} < 0.001$) and 2016 ($PMCM_{A\ vs\ I} = 0.005$, $PMCMC_{A\ vs\ O} = 0.008$). The statistical model assumed a Poisson error distribution and a random effect of the enclosure position on the fixed effect variance.

**Table S1.** Individuals sampled for the *de novo* transcriptome assembly of *I. elegans* used in this study. All individuals come from local populations in Southern Sweden and they were collected either in the field, at their populations of origin, or in experimental outdoor enclosures (Mesocosms), in a field station in the same area. Samples were collected during the mating seasons of 2012 and 2015. The first 27 samples were used for the differential expression analysis in this study.

| ID | Sex | Morph | Colour phase | Season | Collection site | Mating status |
|---|---|---|---|---|---|---|
| T214 | Female | A | immature | 2015 | Field | Unmated |
| T43 | Female | A | immature | 2015 | Field | Unmated |
| T44 | Female | A | immature | 2015 | Field | Unmated |
| T126 | Female | A | transitional | 2015 | Field | Unmated |
| T127 | Female | A | transitional | 2015 | Field | Unmated |
| T45 | Female | A | transitional | 2015 | Field | Unmated |
| T128 | Female | A | mature | 2015 | Field | Unmated |
| T129 | Female | A | mature | 2015 | Field | Unmated |
| T130 | Female | A | mature | 2015 | Field | Unmated |
| T213 | Female | I | immature | 2015 | Field | Unmated |
| T41 | Female | I | immature | 2015 | Field | Unmated |
| T42 | Female | I | immature | 2015 | Field | Unmated |
| 249 | Female | I | transitional | 2015 | Field | Unmated |
| 256 | Female | I | transitional | 2015 | Field | Unmated |
| 266 | Female | I | transitional | 2015 | Field | Unmated |
| T11 | Female | I | mature | 2015 | Field | Mated |
| T29 | Female | I | mature | 2015 | Field | Mated |
| T35 | Female | I | mature | 2015 | Field | Mated |
| T165 | Female | O | immature | 2015 | Field | Unmated |
| T54 | Female | O | immature | 2015 | Field | Unmated |
| T63 | Female | O | immature | 2015 | Field | Unmated |
| T21 | Female | O | transitional | 2015 | Field | Mated |
| T217 | Female | O | transitional | 2015 | Field | Unmated |
| T97 | Female | O | transitional | 2015 | Field | Unmated |
| T96 | Female | O | mature | 2015 | Field | Unmated |
| 188 | Female | O | mature | 2015 | Field | Unmated |
| 248 | Female | O | mature | 2015 | Field | Unmated |
| T234 | Female | I | mature | 2015 | Mesocosm | Unmated |
| T274B7 | Female | I | mature | 2015 | Mesocosm | Mated |
| T267B5-6 | Female | I | mature | 2015 | Mesocosm | Mated |
| T238 | Female | I | mature | 2015 | Mesocosm | Mated |
| T278B7 | Female | I | mature | 2015 | Mesocosm | Mated |
| T242 | Female | I | mature | 2015 | Mesocosm | Mated |
| T250 | Female | I | mature | 2015 | Mesocosm | Unmated |
| T284B7 | Female | I | mature | 2015 | Mesocosm | Unmated |
| T251 | Female | I | mature | 2015 | Mesocosm | Unmated |

| ID | Sex | Morph | Colour phase | Season | Collection site | Mating status |
|---|---|---|---|---|---|---|
| T224 | Female | I | mature | 2015 | Mesocosm | Mated |
| T280B7 | Female | I | mature | 2015 | Mesocosm | Mated |
| T220 | Female | I | mature | 2015 | Mesocosm | Mated |
| T235 | Female | A | mature | 2015 | Mesocosm | Unmated |
| T273B5-6 | Female | A | mature | 2015 | Mesocosm | Unmated |
| T244 | Female | A | mature | 2015 | Mesocosm | Unmated |
| T236 | Female | A | mature | 2015 | Mesocosm | Mated |
| T274B5-6 | Female | A | mature | 2015 | Mesocosm | Mated |
| T273B7 | Female | A | mature | 2015 | Mesocosm | Mated |
| T230 | Female | A | mature | 2015 | Mesocosm | Unmated |
| T280B5-6 | Female | A | mature | 2015 | Mesocosm | Unmated |
| T253 | Female | A | mature | 2015 | Mesocosm | Unmated |
| T222 | Female | A | mature | 2015 | Mesocosm | Mated |
| T284B5-6 | Female | A | mature | 2015 | Mesocosm | Mated |
| T288B7 | Female | A | mature | 2015 | Mesocosm | Mated |
| T172 | Female | I | mature | 2015 | Mesocosm | Unmated |
| T189 | Female | I | mature | 2015 | Mesocosm | Unmated |
| T195 | Female | I | mature | 2015 | Mesocosm | Unmated |
| T171 | Female | A | mature | 2015 | Mesocosm | Unmated |
| T188 | Female | A | mature | 2015 | Mesocosm | Unmated |
| T194 | Female | A | mature | 2015 | Mesocosm | Unmated |
| T70 | Female | A | mature | 2015 | Mesocosm | Mated |
| T81 | Female | A | mature | 2015 | Mesocosm | Mated |
| 50 | Female | A | mature | 2012 | Mesocosm | Unknown |
| 54 | Female | A | mature | 2012 | Mesocosm | Unknown |
| 70 | Female | A | mature | 2012 | Mesocosm | Unknown |
| 80 | Female | A | mature | 2012 | Mesocosm | Unknown |
| 49 | Female | I | mature | 2012 | Mesocosm | Unknown |
| 51 | Female | I | mature | 2012 | Mesocosm | Unknown |
| 52 | Female | I | mature | 2012 | Mesocosm | Unknown |
| 53 | Female | I | mature | 2012 | Mesocosm | Unknown |
| 55 | Female | I | mature | 2012 | Mesocosm | Unknown |
| 56 | Female | I | mature | 2012 | Mesocosm | Unknown |
| 31 | Female | A | mature | 2012 | Mesocosm | Unknown |
| 42 | Female | A | mature | 2012 | Mesocosm | Unknown |
| 43 | Female | A | mature | 2012 | Mesocosm | Unknown |
| 47 | Female | A | mature | 2012 | Mesocosm | Unknown |
| 33 | Female | I | mature | 2012 | Mesocosm | Unknown |
| 35 | Female | I | mature | 2012 | Mesocosm | Unknown |
| 44 | Female | I | mature | 2012 | Mesocosm | Unknown |
| 46 | Female | I | mature | 2012 | Mesocosm | Unknown |

| ID | Sex | Morph | Colour phase | Season | Collection site | Mating status |
|---|---|---|---|---|---|---|
| 8 | Female | A | mature | 2012 | Mesocosm | Unknown |
| 36 | Female | A | mature | 2012 | Mesocosm | Unknown |
| 39 | Female | A | mature | 2012 | Mesocosm | Unknown |
| 40 | Female | A | mature | 2012 | Mesocosm | Unknown |
| 15 | Female | I | mature | 2012 | Mesocosm | Unknown |
| 26 | Female | I | mature | 2012 | Mesocosm | Unknown |
| 38 | Female | I | mature | 2012 | Mesocosm | Unknown |
| 57 | Female | I | mature | 2012 | Mesocosm | Unknown |

**Table S2.** Contrast matrices for the three types of comparisons of differential gene expression in females of *Ischnura elegans*, given the interaction term in the model formula: ~0 + Morph + Colour Phase + Morph: Colour Phase. There are three morphs (Androchrome, Infuscans, and Obsoleta) and three colour development phases: *immature* (Imm), *transitional* (Tra) and *mature* (Mat). The *immature* colour phase is treated as the intercept within each morph (i.e. 'Androchrome' refers to *immature* Androchrome females). Three types of contrasts were estimated: (1) contrasts among morphs within each colour phase, (2) contrasts between subsequent colour phases within morphs and (3) and contrasts between developmental changes among morphs. For (3), developmental changes corresponded to one of two developmental windows: *early* colour development (E), between the *immature* and *transitional* phase and *late* colour development (L), between the *transitional* and *mature* colour phase.

**(1) Among morphs** — **Contrasts**

| Levels | AImmvsIImm | AImmvsOImm | IImmvsOImm | ATravsITra | ATravsOTra | ITravsOTra | AMatvsIMat | AMatvsOMat | IMatvsOMat |
|---|---|---|---|---|---|---|---|---|---|
| Androchrome | 1 | 1 | 0 | 0 | 0 | 0 | 0 | 0 | 0 |
| Infuscans | -1 | 0 | 1 | 0 | 0 | 0 | 0 | 0 | 0 |
| Obsoleta | 0 | -1 | -1 | 0 | 0 | 0 | 0 | 0 | 0 |
| AgeTransitional | 0 | 0 | 0 | 1 | 1 | 0 | 0 | 0 | 0 |
| Infuscans.AgeTransitional | 0 | 0 | 0 | -1 | 0 | 1 | 0 | 0 | 0 |
| Obsoleta.AgeTransitional | 0 | 0 | 0 | 0 | -1 | -1 | 0 | 0 | 0 |
| AgeMature | 0 | 0 | 0 | 0 | 0 | 0 | 1 | 1 | 0 |
| Infuscans.AgeMature | 0 | 0 | 0 | 0 | 0 | 0 | -1 | 0 | 1 |
| Obsoleta.AgeMature | 0 | 0 | 0 | 0 | 0 | 0 | 0 | -1 | -1 |

**(2) Between phases** — **Contrasts**

| Levels | ATravsAImm | ITravsIImm | OTravsOImm | AMatvsATra | IMatvsITra | OMatvsOTra |
|---|---|---|---|---|---|---|
| Androchrome | -1 | 0 | 0 | 0 | 0 | 0 |
| Infuscans | 0 | -1 | 0 | 0 | 0 | 0 |
| Obsoleta | 0 | 0 | -1 | 0 | 0 | 0 |
| AgeTransitional | 1 | 0 | 0 | -1 | 0 | 0 |
| Infuscans.AgeTransitional | 0 | 1 | 0 | 0 | -1 | 0 |
| Obsoleta.AgeTransitional | 0 | 0 | 1 | 0 | 0 | -1 |

| | | | | | | |
|---|---|---|---|---|---|---|
| AgeMature | 0 | 0 | 0 | 1 | 0 | 0 |
| Infuscans.AgeMature | 0 | 0 | 0 | 0 | 1 | 0 |
| Obsoleta.AgeMature | 0 | 0 | 0 | 0 | 0 | 1 |

| **(3) Phase differences among morphs** | | | **Contrasts** | | | |
|---|---|---|---|---|---|---|
| Levels | AEvsIE | AEvsOE | IEvsOE | ALvsIL | ALvsOL | ILvsOL |
| Androchrome | -1 | -1 | 0 | 0 | 0 | 0 |
| Infuscans | 1 | 0 | -1 | 0 | 0 | 0 |
| Obsoleta | 0 | 1 | 1 | 0 | 0 | 0 |
| AgeTransitional | 1 | 1 | 0 | -1 | -1 | 0 |
| Infuscans.AgeTransitional | -1 | 0 | 1 | 1 | 0 | -1 |
| Obsoleta.AgeTransitional | 0 | -1 | -1 | 0 | 1 | 1 |
| AgeMature | 0 | 0 | 0 | 1 | 1 | 0 |
| Infuscans.AgeMature | 0 | 0 | 0 | -1 | 0 | 1 |
| Obsoleta.AgeMature | 0 | 0 | 0 | 0 | -1 | -1 |

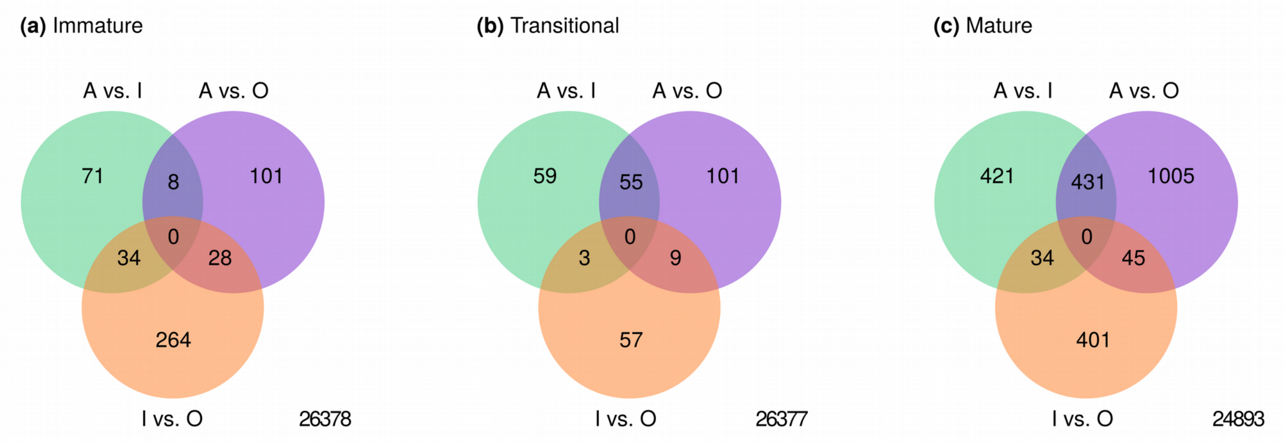


**Figure S2.** Venn diagram of differentially expressed genes between morphs and within developmental colour phase in females of *I. elegan*s (contrast group (1); Table S2). Three females of each morph were sampled at each of three stages of adult colour development **(a)** *immature*, **(b)** *transitional* and **(c)** *mature*. The number of genes with no evidence of morph differences in differential expression between developmental colour phases is shown in the bottom right corner of each plot.

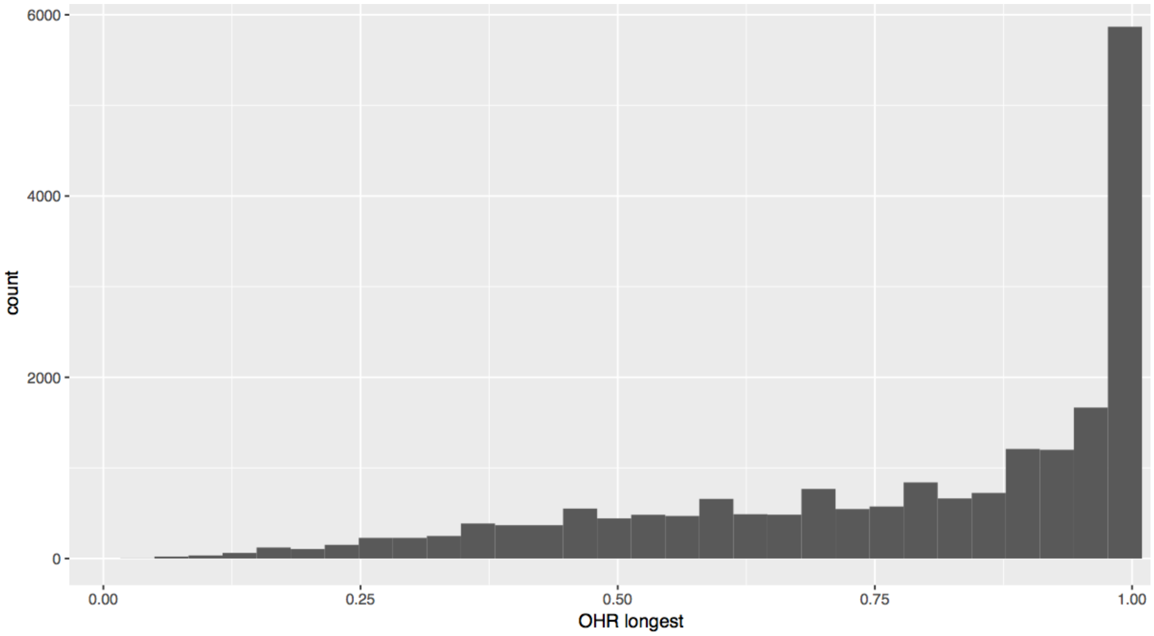


**Figure S3.** Histogram of ortholog hit ratios (OHR) for *Ischnura elegans* transcripts with orthologs in the *Calopteryx splendens* genome. The OHR represents the fraction of the protein coding sequences in the *C. splendens* genome that is covered by a single (longest) *I. elegans* transcript. Around 70% of the sequences are covered to 70% or more of their full expected length, indicating the transcriptome quality is acceptable.

**Table S3.** Distribution of protein clusters in the *Calopteryx splendens* genome with orthologs in the *Ischnura elegans* transctriptome. The first column indicates the sequence length in the *C. splendens* genome and the remaining columns show the ortholog hit ratio (OHR) of the longest transcript in the *I. elegans* transcriptome mapped to each *C. splendens* protein coding sequence. The OHR represents the fraction of the protein coding sequence that is covered by a single (longest) *I. elegans* transcript. Around 70% of the sequences are covered to 70% or more of their full expected length, indicating the transcriptome quality is acceptable.

| | Assembly length quantile | | | | | | | | | |
|---|---|---|---|---|---|---|---|---|---|---|
| **Length** | **0-0.1** | **0.11-0.2** | **0.21-0.3** | **0.31-0.4** | **0.41-0.5** | **0.51-0.6** | **0.61-0.7** | **0.71-0.8** | **0.81-0.9** | **0.9-1.0** |
| 0-100 | 0 | 0 | 6 | 44 | 116 | 155 | 201 | 265 | 320 | 1033 |
| 101-200 | 0 | 21 | 119 | 233 | 337 | 410 | 484 | 629 | 866 | 3213 |
| 201-500 | 13 | 127 | 176 | 226 | 261 | 322 | 412 | 564 | 940 | 4858 |
| 501-1000 | 12 | 30 | 21 | 29 | 38 | 56 | 105 | 481 | 374 | 1938 |
| 1001-2000 | 2 | 7 | 3 | 9 | 8 | 8 | 15 | 46 | 117 | 491 |
| 2001-5000 | 0 | 0 | 0 | 3 | 1 | 1 | 03 | 5 | 15 | 85 |
| 5001-10000 | 0 | 0 | 0 | 0 | 0 | 1 | 0 | 0 | 2 | 6 |
| >10000 | 0 | 0 | 0 | 0 | 0 | 0 | 0 | 0 | 0 | 0 |
| Sum | 27 | 185 | 325 | 544 | 761 | 953 | 1220 | 1689 | 2634 | 11624 |

**Table S4.** GO terms for significantly enriched biological processes (Fisher's P < 0.01) in the subset of downregulated genes shared by all female morphs of of *I. elegans*, during *early* colour development.

| Rank | GO ID | Term | Annotated | Significant | Expected | Fisher's P |
|---|---|---|---|---|---|---|
| 1 | GO:0006412 | translation | 619 | 214 | 116.94 | 7.5e-18 |
| 2 | GO:0015991 | ATP hydrolysis coupled proton transport | 44 | 21 | 8.31 | 1.3e-5 |
| 3 | GO:0015986 | ATP synthesis coupled proton transport | 35 | 18 | 6.61 | 1.4e-5 |
| 4 | GO:0006886 | intracellular protein transport | 237 | 71 | 44.77 | 2.0e-5 |
| 5 | GO:1902600 | proton transmembrane transport | 115 | 55 | 21.73 | 6.5e-5 |
| 6 | GO:0006511 | ubiquitin-dependent protein catabolic process | 119 | 38 | 22.48 | 4.3e-4 |
| 7 | GO:0007264 | small GTPase mediated signal transduction | 215 | 60 | 40.62 | 7.1e-4 |
| 8 | GO:0001731 | formation of translation preinitiation complex | 16 | 9 | 3.02 | 9.5e-4 |
| 9 | GO:0006414 | translational elongation | 65 | 23 | 12.28 | 0.001 |
| 10 | GO:0042423 | catecholamine biosynthetic process | 4 | 4 | 0.76 | 0.001 |
| 11 | GO:0016579 | protein deubiquitination | 47 | 18 | 8.88 | 0.001 |
| 12 | GO:0009108 | coenzyme biosynthetic process | 120 | 36 | 22.67 | 0.002 |
| 13 | GO:0048193 | Golgi vesicle transport | 43 | 16 | 8.12 | 0.004 |
| 14 | GO:0045116 | protein neddylation | 5 | 4 | 0.94 | 0.005 |
| 15 | GO:0006465 | signal peptide processing | 5 | 4 | 0.94 | 0.005 |
| 16 | GO:0042773 | ATP synthesis coupled electron transport | 23 | 10 | 4.35 | 0.006 |
| 17 | GO:0051603 | proteolysis involved in cellular protein catabolic process | 139 | 47 | 26.26 | 0.006 |
| 18 | GO:0006413 | translational initiation | 140 | 44 | 26.45 | 0.007 |
| 19 | GO:0019243 | methylglyoxal catabolic process to D-lactate via S-lactoyl-glutathione | 3 | 3 | 0.57 | 0.007 |
| 20 | GO:0006826 | iron ion transport | 14 | 7 | 2.64 | 0.008 |
| 21 | GO:0035194 | posttranscriptional gene silencing by RNA | 11 | 6 | 2.08 | 0.009 |
| 22 | GO:0006163 | purine nucleotide metabolic process | 208 | 68 | 39.3 | 0.010 |

**Table S5.** GO terms for significantly enriched biological processes (Fisher's P < 0.01) in the subset of upregulated genes for each female morph of of *I. elegans*, during *early* colour development. GO terms with fewer than 3 annotated genes are excluded.

| Rank | GO ID | Term | Annotated | Significant | Expected | Fisher's P |
|---|---|---|---|---|---|---|
| *A-females* | | | | | | |
| 1 | GO:0090304 | nucleic acid metabolic process | 1832 | 16 | 7.77 | 0.002 |
| *I-females* | | | | | | |
| 1 | GO:0006278 | RNA-dependent DNA biosynthetic process | 206 | 7 | 1.08 | 8.4e-5 |
| 2 | GO:1901136 | carbohydrate derivative catabolic process | 60 | 3 | 0.31 | 0.004 |
| *O-females* | | | | | | |
| 1 | GO:0071555 | cell wall organization | 21 | 5 | 0.55 | 1.7e-4 |
| 2 | GO:0005980 | glycogen catabolic process | 5 | 3 | 0.13 | 1.7e-4 |
| 3 | GO:0072488 | ammonium transmembrane transport | 6 | 3 | 0.16 | 3.3-4 |
| 4 | GO:0006355 | regulation of transcription, DNA-templated | 593 | 29 | 15.54 | 7.6e-4 |
| 5 | GO:0007094 | mitotic spindle assembly checkpoint | 9 | 3 | 0.24 | 0.001 |
| 6 | GO:0055123 | digestive system development | 5 | 2 | 0.13 | 0.006 |
| 7 | GO:0045165 | cell fate commitment | 6 | 2 | 0.16 | 0.010 |

**Table S6.** GO terms for significantly enriched biological processes (Fisher's P < 0.01) in the subset of downregulated genes for each female morph of of *I. elegans*, during *late* colour development. GO terms with fewer than 3 annotated genes are excluded.

| Rank | GO ID | Term | Annotated | Significant | Expected | Fisher's P |
|---|---|---|---|---|---|---|
| *A-females* | | | | | | |
| 1 | GO:0015986 | ATP synthesis coupled proton transport | 35 | 21 | 1.92 | 2.4e-18 |
| 2 | GO:0055114 | oxidation-reduction process | 836 | 129 | 45.9 | 5.6e-17 |
| 3 | GO:1902600 | proton transmembrane transport | 115 | 45 | 6.31 | 1.8e-16 |
| 4 | GO:0022900 | electron transport chain | 36 | 26 | 1.98 | 1.4e-7 |
| 5 | GO:0006120 | mitochondrial electron transport, NADH to ubiquinone | 7 | 6 | 0.38 | 1.8e-7 |
| 6 | GO:0042773 | ATP synthesis coupled electron transport | 23 | 16 | 1.26 | 1.7e-6 |
| 7 | GO:0006099 | tricarboxylic acid cycle | 30 | 9 | 1.65 | 2.1e-5 |
| 8 | GO:0006108 | malate metabolic process | 11 | 5 | 0.6 | 1.7e-4 |
| 9 | GO:0042775 | mitochondrial ATP synthesis coupled electron transport | 14 | 10 | 0.77 | 2.6e-4 |
| 10 | GO:0006744 | ubiquinone biosynthetic process | 8 | 4 | 0.44 | 5.3e-4 |
| 11 | GO:0034220 | ion transmembrane transport | 347 | 73 | 19.05 | 6.6e-4 |
| 12 | GO:0070588 | calcium ion transmembrane transport | 22 | 6 | 1.21 | 9.3e-4 |
| 13 | GO:0006777 | Mo-molybdopterin cofactor biosynthetic process | 10 | 4 | 0.55 | 0.001 |
| 14 | GO:0019646 | aerobic electron transport chain | 5 | 3 | 0.27 | 0.002 |
| 15 | GO:0016180 | snRNA processing | 11 | 4 | 0.6 | 0.002 |
| 16 | GO:0015991 | ATP hydrolysis coupled proton transport | 44 | 8 | 2.42 | 0.002 |
| 17 | GO:0022904 | respiratory electron transport chain | 29 | 20 | 1.59 | 0.008 |
| *I-females* | | | | | | |
| 1 | GO:0006633 | fatty acid biosynthetic process | 55 | 8 | 0.87 | 2.1e-6 |
| 2 | GO:0006869 | lipid transport | 75 | 9 | 1.19 | 2.5e-6 |
| 3 | GO:0035194 | posttranscriptional gene silencing by RNA | 11 | 3 | 0.17 | 5.8e-4 |
| 4 | GO:0032259 | methylation | 224 | 10 | 3.55 | 0.003 |
| 5 | GO:0006629 | lipid metabolic process | 281 | 17 | 4.45 | 0.007 |
| 6 | GO:0006026 | aminoglycan catabolic process | 48 | 4 | 0.76 | 0.007 |
| *O-females* | | | | | | |
| 1 | GO:0035335 | peptidyl-tyrosine dephosphorylation | 46 | 5 | 0.75 | 8.4e-4 |
| 2 | GO:0009157 | deoxyribonucleoside monophosphate biosynthetic process | 4 | 2 | 0.06 | 0.002 |
| 3 | GO:0007062 | sister chromatid cohesion | 18 | 3 | 0.29 | 0.003 |
| 4 | GO:0044772 | mitotic cell cycle phase transition | 38 | 4 | 0.62 | 0.003 |
| 5 | GO:0009451 | RNA modification | 81 | 5 | 1.31 | 0.010 |

**Table S7.** GO terms for significantly enriched biological processes (Fisher's P < 0.01) in the subset of upregulated genes for each female morph of of *I. elegans*, during *late* colour development. GO terms with fewer than 3 annotated genes are excluded.

| Rank | GO ID | Term | Annotated | Significant | Expected | Fisher's P |
|---|---|---|---|---|---|---|
| *A-females* | | | | | | |
| 1 | GO:0006260 | DNA replication | 98 | 40 | 15.07 | 1.5e-5 |
| 2 | GO:0034587 | piRNA metabolic process | 5 | 5 | 0.77 | 8.5e-5 |
| 3 | GO:0071139 | resolution of recombination intermediates | 5 | 5 | 0.77 | 8.5e-5 |
| 4 | GO:0006468 | protein phosphorylation | 452 | 99 | 69.49 | 9.5e-5 |
| 5 | GO:0006281 | DNA repair | 167 | 49 | 25.67 | 1.3e-4 |
| 6 | GO:0006261 | DNA-dependent DNA replication | 36 | 15 | 5.53 | 1.3e-4 |
| 7 | GO:0000724 | double-strand break repair via homologous recombination | 15 | 8 | 2.31 | 7.1e-4 |
| 8 | GO:0007049 | cell cycle | 190 | 64 | 29.21 | 8.2e-4 |
| 9 | GO:0051304 | chromosome separation | 16 | 8 | 2.46 | 0.001 |
| 10 | GO:0007076 | mitotic chromosome condensation | 7 | 5 | 1.08 | 0.001 |
| 11 | GO:0090501 | RNA phosphodiester bond hydrolysis | 49 | 16 | 7.53 | 0.002 |
| 12 | GO:0051276 | chromosome organization | 227 | 66 | 34.9 | 0.002 |
| 13 | GO:0034314 | Arp2/3 complex-mediated actin nucleation | 17 | 8 | 2.61 | 0.002 |
| 14 | GO:0007062 | sister chromatid cohesion | 18 | 8 | 2.77 | 0.003 |
| 15 | GO:0007346 | regulation of mitotic cell cycle | 48 | 15 | 7.38 | 0.004 |
| 16 | GO:0051301 | cell division | 62 | 18 | 9.53 | 0.004 |
| 17 | GO:0097659 | nucleic acid-templated transcription | 718 | 142 | 110.38 | 0.006 |
| 18 | GO:0000075 | cell cycle checkpoint | 36 | 12 | 5.53 | 0.006 |
| 19 | GO:0016570 | histone modification | 78 | 21 | 11.99 | 0.006 |
| 20 | GO:0006869 | lipid transport | 75 | 20 | 11.53 | 0.008 |
| 21 | GO:0006355 | regulation of transcription, DNA-templated | 593 | 112 | 91.16 | 0.009 |
| 22 | GO:0007017 | microtubule-based process | 159 | 36 | 24.44 | 0.009 |
| 23 | GO:0051321 | meiotic cell cycle | 17 | 7 | 2.61 | 0.009 |
| *I-females* | | | | | | |
| 1 | GO:1902600 | proton transmembrane transport | 115 | 10 | 1.05 | 7.3e-8 |
| 2 | GO:0006119 | oxidative phosphorylation | 38 | 5 | 0.35 | 2.2e-5 |
| 3 | GO:0015893 | drug transport | 14 | 3 | 0.13 | 2.5e-4 |
| 4 | GO:0019646 | aerobic electron transport chain | 5 | 2 | 0.05 | 8.0e-4 |
| 5 | GO:0035725 | sodium ion transmembrane transport | 22 | 3 | 0.2 | 9.9e-4 |
| 6 | GO:0045333 | cellular respiration | 72 | 8 | 0.66 | 0.002 |
| 7 | GO:0022900 | electron transport chain | 36 | 5 | 0.33 | 0.003 |
| 8 | GO:0006885 | regulation of pH | 11 | 2 | 0.1 | 0.004 |
| 9 | GO:0009060 | aerobic respiration | 45 | 5 | 0.41 | 0.005 |
| 10 | GO:0006754 | ATP biosynthetic process | 89 | 4 | 0.81 | 0.009 |
| O-females | | | | | | |
| 1 | GO:0022900 | electron transport chain | 36 | 6 | 0.22 | 4.9e-5 |
| 2 | GO:0042773 | ATP synthesis coupled electron transport | 23 | 3 | 0.14 | 3.5e-4 |

| Rank | GO ID | Term | Annotated | Significant | Expected | Fisher’s P |
|---|---|---|---|---|---|---|
| 3 | GO:0055114 | oxidation-reduction process | 836 | 16 | 5.11 | 0.009 |

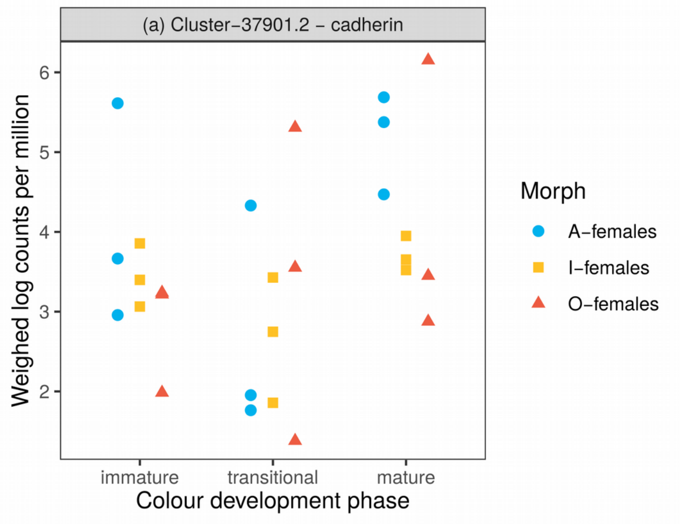


**Figure S4.** Expression pattern of gene annotated to cadherin, which was co-regulated and potentially in epistasis (see Discussion) with a heterogeneous ribonucleoprotein during adult development of *I. elegans* females. Transcripts were annotated against the non-redundant database of the National Center for Biotechnology Information (NCBI). Expression data (log counts per million) have been weighed by within-sample variability and by the mean-variance relationship among genes, using ‘voom’ (see Methods).

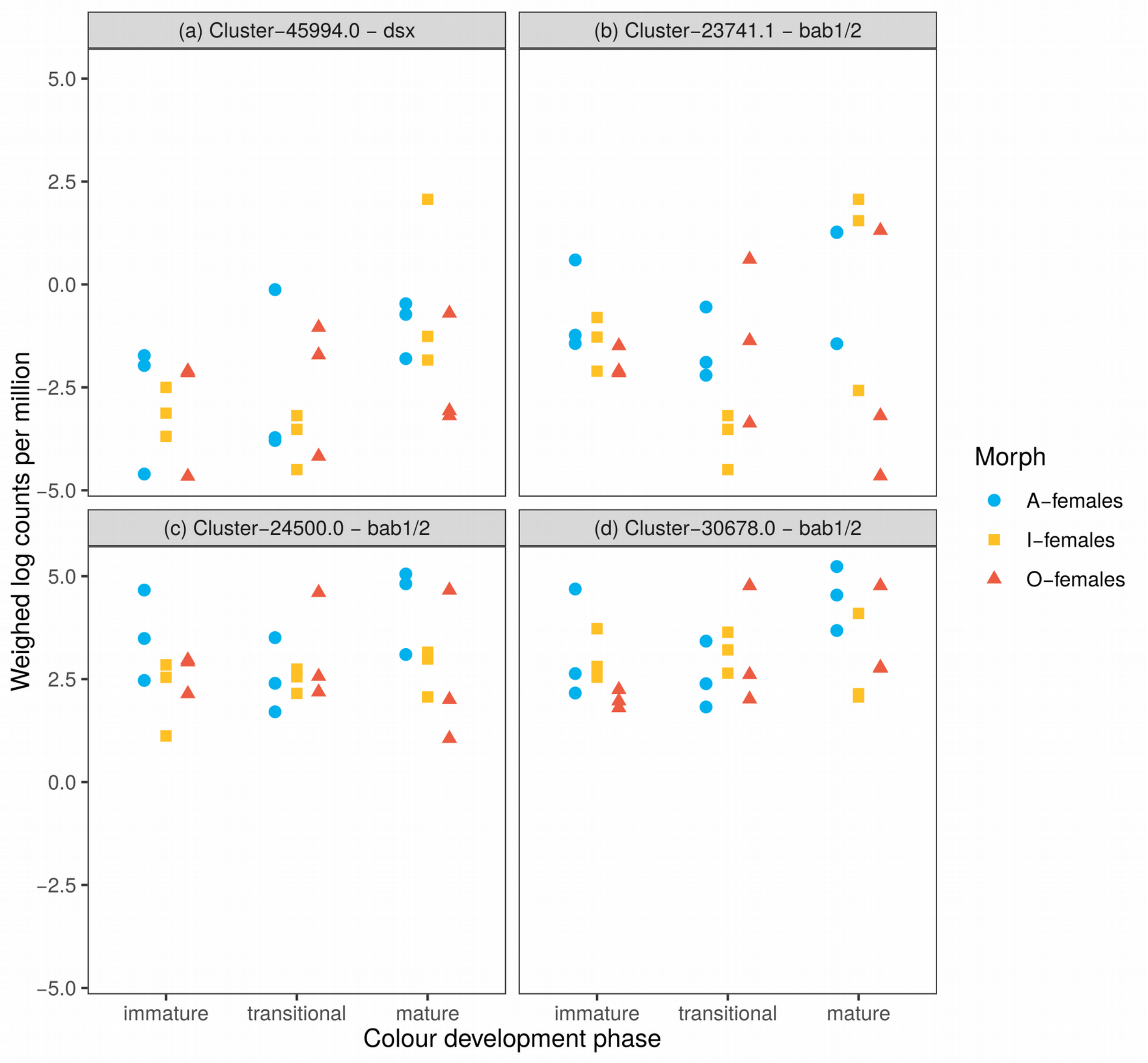


**Figure S5.** Expression patterns of genes in the *I. elegans* transcriptome with significant blast hits against *doublesex* (*dsx*) and on of its direct targets, bric-á-brac 1 and 2 (bab1/2) in *D. melanogaster.* Transcripts were annotated against the non-redundant database of the National Center for Biotechnology Information (NCBI). Expression data (log counts per million) have been weighed by within-sample variability and by the mean-variance relationship among genes, using 'voom' (see Methods).
